# The genetic architecture of an allosteric hormone receptor

**DOI:** 10.1101/2025.05.30.656975

**Authors:** Maximilian R. Stammnitz, Ben Lehner

## Abstract

Many proteins function as switches, detecting chemicals and transducing their concentrations into cellular responses. Receptor switches are key to the integration of environmental signals, yet it is not well understood how their signal processing is genetically encoded. Here, to address these questions, we present a complete map of how mutations quantitatively alter the input-output function of a receptor. Using a massively parallel approach, GluePCA, we quantify more than 40,000 binding measurements and 3,500 dose-response curves for variants of the allosteric phytohormone receptor PYL1. Strikingly, >91% of missense variants tune the dose-response of the receptor. Many variants drive changes in sensitivity, basal activity, maximum response and induction steepness in a correlated manner, suggesting a shared underlying biophysical mechanism, which we identify as stability changes. Beyond this, signalling parameters can be independently tuned by mutations, with surprisingly large effects in interface-distal positions and a modular genetic architecture across the receptor’s structure. A small subset of missense variants confers qualitative phenotypic innovation, producing hypersensitive and inverted response patterns. Our data demonstrate the feasibility of dose-response profile quantification at massive scale and reveal the remarkable evolutionary malleability of an allosteric switch.

## Introduction

The human genome encodes more than 1,000 receptors, proteins that bind to specific ligands such as hormones, neurotransmitters, antigens and other small molecules to elicit cellular responses (*1, 2*). Receptors are often located on the cell surface but can also reside in the cytoplasm and nucleus. Important examples of intracellular receptors include mammalian steroid hormone receptors and many plant hormone sensors (*3, 4*). Receptors work as molecular switches, converting extracellular or intracellular signals into a biological effect. Despite their structural diversity, most receptors share a common mechanism of action: the binding of a ligand to one region of the protein, which triggers a change in the conformation or dynamics of the peptide chain to initiate a response at a different region of the protein. This long-range transmission of information in proteins is termed allostery, which, due to its central role in regulation, has been referred to as ‘the second secret of life’ (*5, 6*). Allostery allows receptors to function as tunable molecular relays, finely altering their activity in response to changes in ligand concentration (*7*).

How the allosteric input-output functions of receptors are encoded in their amino acid sequences is largely unexplored. Receptor activity often follows a sigmoidal (S-shaped) dose-response curve, characterised by a basal response at low ligand concentration, a steep increase over intermediate concentrations, and a saturated maximal response at high concentrations. The sigmoidal dose-response curves of many receptors can be well-approximated using Hill functions with four parameters: basal activity (*B*_*0*_), maximum activity (*B*_*∞*_), sensitivity as determined by the concentration at half-maximum response (*EC*_*50*_), and the steepness of the response (Hill coefficient, *n*) (Fig. 1A). Theoretical models of allosteric proteins predict that changes in many different biophysical properties can affect the four Hill function parameters (*8, 9*). The introduction of random combinations of amino acid substitutions in allosteric prokaryotic transcription factors has revealed that both individual and multi-mutants frequently alter their dose-response curves (*10, 11*). Similarly, individually quantifying the Hill curves of an allosteric enzyme (*12*) and G protein-coupled receptors (*13*–*15*) with residues mutated to alanine or glycine has revealed diverse and frequent consequences. However, this latter approach of characterising mutated receptors one-by-one would be difficult to scale to the size of an experiment required to characterise the effects of all mutations in all positions in a protein.

**Figure 1:**
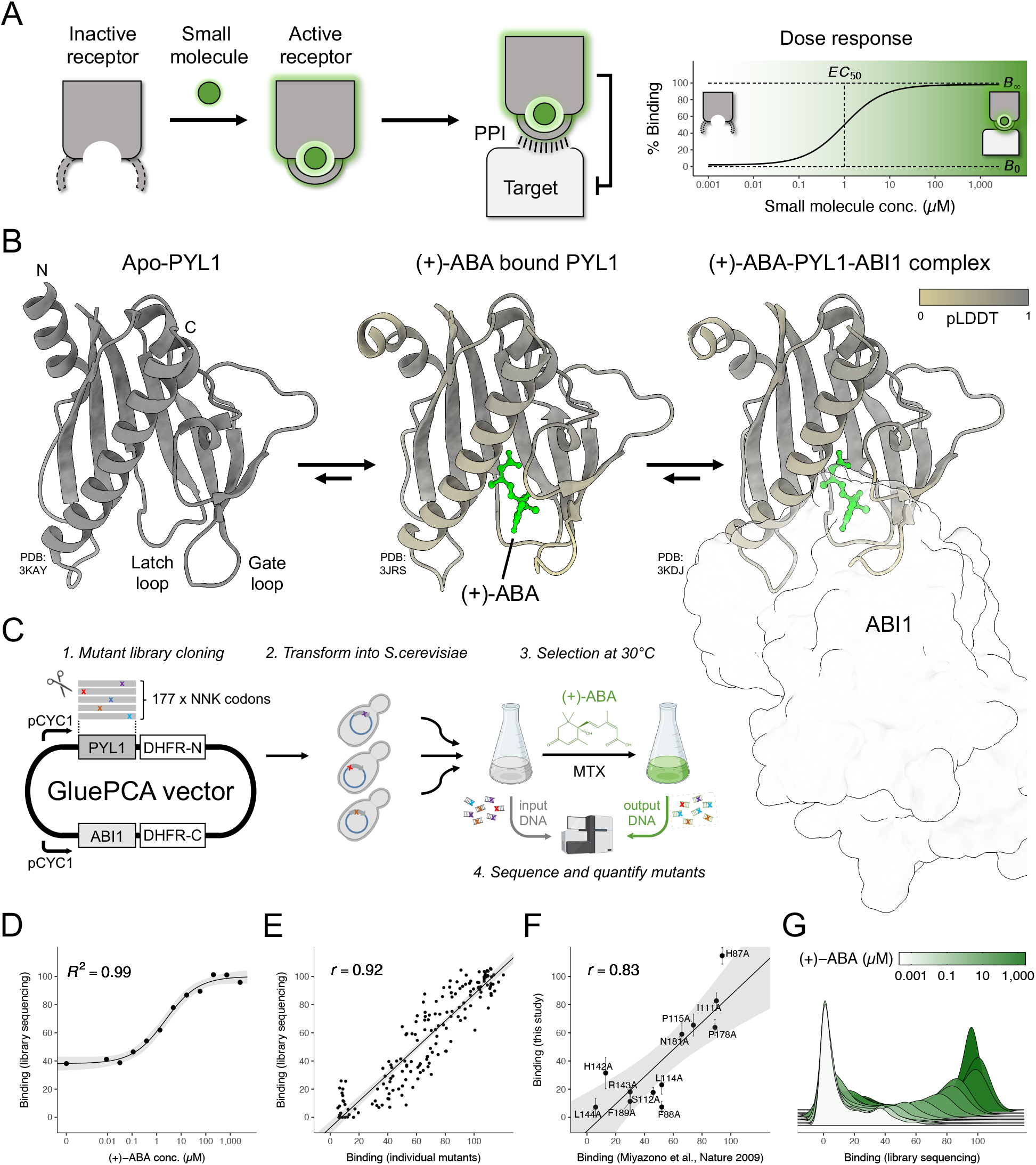
Deep mutational scanning of inducible PYL1 receptor binding by GluePCA. **A:** Schematics of small molecule ligand-induced activation of an allosteric receptor protein. As in the case of PYL1, target inhibition is well approximated by a log-logistic response of ligand concentration. **B:** PYL1 structural changes upon ligand binding. Left: Apo-PYL1 crystal structure displayed as a ribbon diagram. Centre: (+)-ABA bound PYL1 structure, with the (+)-ABA molecule highlighted in green. Right: (+)-ABA-PYL1-ABI1 complex, in which the ABI1 phosphatase silhouette is set to high transparency. PYL1 residue colouring in the (+)-ABA bound structures is scaled by predicted local distance difference test (pLDDT) with respect to the Apo-PYL1 structure, highlighting dynamic positions. **C:** GluePCA setup for deep mutational scanning measurements of inducible protein-protein binding. On the same plasmid, two CYC1 promoters express PYL1 mutants as N-terminal fusions to DHFR-N and wildtype ABI1 as an N- terminal fusion to DHFR-C. Upon variant library transformation into *S*.*cerevisiae*, changes in relative cell fractions are obtained through a series of variant competition experiments in which vital DHFR reconstitution is mediated through (+)-ABA dependent PYL1-ABI1 association. Mutant-specific fitness at each (+)-ABA concentration is calculated from deep sequencing and relative PYL1 variant counting prior and after selection (see Materials & Methods). **D:** Wildtype PYL1-ABI1 binding dose response regression from a single twelve-step (+)-ABA titration. Data points represent bulk library GluePCA fitness estimates from deep sequencing, line four-parametric Hill model regression, grey shading 95% confidence interval. *R*^*2*^, coefficient of determination (see Supplementary Fig. S2A-B). **E:** ABI1-binding of 14 PYL1 variants at twelve (+)-ABA concentrations, as measured by bulk library GluePCA selection and deep sequencing versus separate, single-variant GluePCA growth measurements. Line represents linear regression (*P* < 1×10^−15^), grey shading 95% confidence interval. *r*, Pearson correlation coefficient. **F:** ABI1-binding of twelve alanine mutants of PYL1, as measured by bulk library GluePCA versus *in vitro* GST pull- down assays (10 µM (+)-ABA), respectively scaled by wildtype PYL1-ABI1 association strength. Line represents linear regression (*P* < 1×10^−4^), grey shading 95% confidence interval. *r*, Pearson correlation coefficient (see Supplementary Fig. S2C). **G:** Kernel density of bulk PYL1 mutant library GluePCA binding fitness at varying (+)-ABA concentrations. Green colour coding indicates increasing hormone level from 0 nM (brightest) to 2,500 µM (darkest) (see Supplementary Fig. S2D).

Even the simplest cellular networks typically elicit non-linear responses and contain one or more feedback loops. This makes it challenging to disentangle change in the intrinsic input-output function of a receptor from measurement of a downstream response (*16*). To characterise the genetic architectures of receptors it is therefore highly desirable to measure the molecular activity of a receptor as directly and proximally as possible. One class of receptors where such proximal measurements are possible is plant hormone receptors. Similar to mammalian hormone receptors, phytohormone sensors such as the strigolactone, gibberellin and abscisic acid receptors bind to their small molecule ligands and undergo conformational changes that alter their affinity to other macromolecules (*3, 17, 18*). This chemically-induced dimerisation (*18*–*21*) can be readily quantified using protein-protein interaction assays in non-plant cell systems and *in vitro*, allowing direct measurement of the proximal molecular response to ligand engagement. For example, binding of the stress hormone abscisic acid ((+)-ABA) to Pyrabactin Resistance 1-Like (PYR/PYL) family receptors triggers a major conformational change in two loops, termed ‘gate’ and ‘latch’, that allows binding to class 2C protein phosphatases (PP2C) such as ABA-Insensitive 1 (ABI1) (Fig. 1B). This complex formation inhibits the PP2Cs, which in turn leads to the expression of drought-response genes – a highly conserved molecular signalling pathway with central relevance to global food security (*22, 23*).

The main structural changes in PYR/PYL proteins upon hormone binding are well characterised (*24*–*26*), making them an attractive model system for studying deep receptor genotype-phenotype maps. PYR/PYL structures belong to the conserved START domain superfamily of helix-grip folds, which form central hydrophobic cavities that are evolutionarily co-opted to fit diverse ligands (*24*–*30*). Members of the PYR/PYL receptor family feature a substantial diversity in ability to sense (+)-ABA and to inhibit different phosphatases. This has been primarily attributed to a select few residues in the hormone pocket and protein-protein interface (PPI), which can also be used to reprogram the receptors’ ligand specificity towards structurally diverse small-molecules via mutations (*31*–*39*). These interfaces allow the receptors to be repurposed for biotechnological applications, including as sensors for pharmacological and environmental chemicals and for agrochemical control of plant resistance under water stress (*34*–*47*). Beyond these focal sites, however, the allosteric nature of chemically-induced PYR/PYL remodelling remains obscure.

Here we present a massively parallel, pooled experimental approach that allows us to quantify receptor dose-response curves at unprecedented scale and apply it to understand the genetic architecture of the PYR/PYL switch.

## Results

### GluePCA: massively parallel quantification of chemically-induced dimerisation

To measure changes in the activity of intracellular receptors at scale, we developed a high-throughput method that quantifies chemically induced dimerization (CID) using massively parallel DNA synthesis, *in vivo* selection and deep next-generation sequencing. The approach, which we term Glueable Protein-fragment Complementation Assay (GluePCA) fuses a receptor and target protein to two different fragments of an essential enzyme, dihydrofolate reductase (DHFR). Addition of a chemical that induces association between receptor and target reconstitutes DHFR and thereby allows for growth of yeast cells under selective conditions (Fig. 1C). The scalability and dynamic range of DHFR-based protein complementation assays has been benchmarked for interactions of thousands of native yeast proteins (*48, 49*), hundreds of human proteins (*50*) and millions of mutants thereof (*50*–*56*).

To establish the generality of GluePCA, we first tested its ability to detect ligand-induced interactions for a range of structurally diverse proteins. Characteristic chemical dimerization dose-response curves were detectable for natural CID systems, including PYL1-ABI1 mediated by (+)- ABA, GID1A-GAI by gibberellic acid (GA3), and FKBP12-mTOR/FRB by rapamycin (Supplementary Fig. S1). Testing the assay’s dynamic range through previously reported CID- disrupting mutations in ABI1 (W300A) (*24*), GAI (E54R) (*57*), and mTOR/FRB (PLF Y2088A) (*58, 59*) suggests that GluePCA is a suitable growth-based selection assay to quantify small molecule-induced protein-protein interactions. This allows for pooled quantification of chemically induced dimerization of different genotypes in a single high-throughput experiment (Supplementary Fig. S1).

### Quantifying >3,500 dose-response curves

We used degenerate oligonucleotide synthesis to construct a library containing all 3,363 single amino acid substitutions, 177 stop codons and 214 synonymous wildtype variants in *Arabidopsis thaliana* PYL1 (residues 33-209 (*40*)) fused to an N-terminal fragment of DHFR (Supplementary Table S1). The library was then transformed into yeast and we quantified its ability to bind to ABI1 (residues 126-423 (*40*)) fused to the DHFR C-terminus. This was done in pooled, plasmid-based selection experiments across 12 different ABA concentrations (*60*). Our multiscale CID binding measurements are highly reproducible (*r* = 0.92, N = 168; Fig. 1D-E, Supplementary Fig. S2A-B) and also very well correlated with independent *in vitro* measurements (*r* = 0.83, N = 12; Fig. 1F, Supplementary Fig. S2C), further validating the approach.

### The genetic landscape of a hormone receptor

In total we made 42,301 binding measurements (3,511-3,529 mutations (99.2-99.7% programmed) at 12 (+)-ABA concentrations (Supplementary Table S2). Plotting the distribution of binding scores at each ligand concentration (Fig. 1G) reveals a bimodal distribution of binding at most (+)-ABA concentrations, with the binding strength of the upper wildtype-like mode increasing as the concentration of hormone is increased. In the absence of ligand, >85% of variants have no detectable binding to ABI1 (Fig. 2A, Supplementary Fig. S2D). As the ligand concentration is increased, a larger percentage of variants show wildtype-like binding, consistent with binding-induced protein stabilisation (*61, 62*). At the highest ligand concentration of 2,500 µM (+)-ABA, >85% of variants have binding similar to wildtype PYL1 (Fig. 2A, Supplementary Fig. S2D). The most detrimental mutations are strongly enriched in the protein core (Fisher’s exact test, *P* < 10^− 15^, odds ratio (*OR*) = 7.6), as well as in (+)-ABA (*P* < 10^−15^, *OR* = 6.5) and ABI1-contacting (*P* < 10^−8^, *OR* = 1.8) residues (Supplementary Fig. S2E).

**Figure 2:**
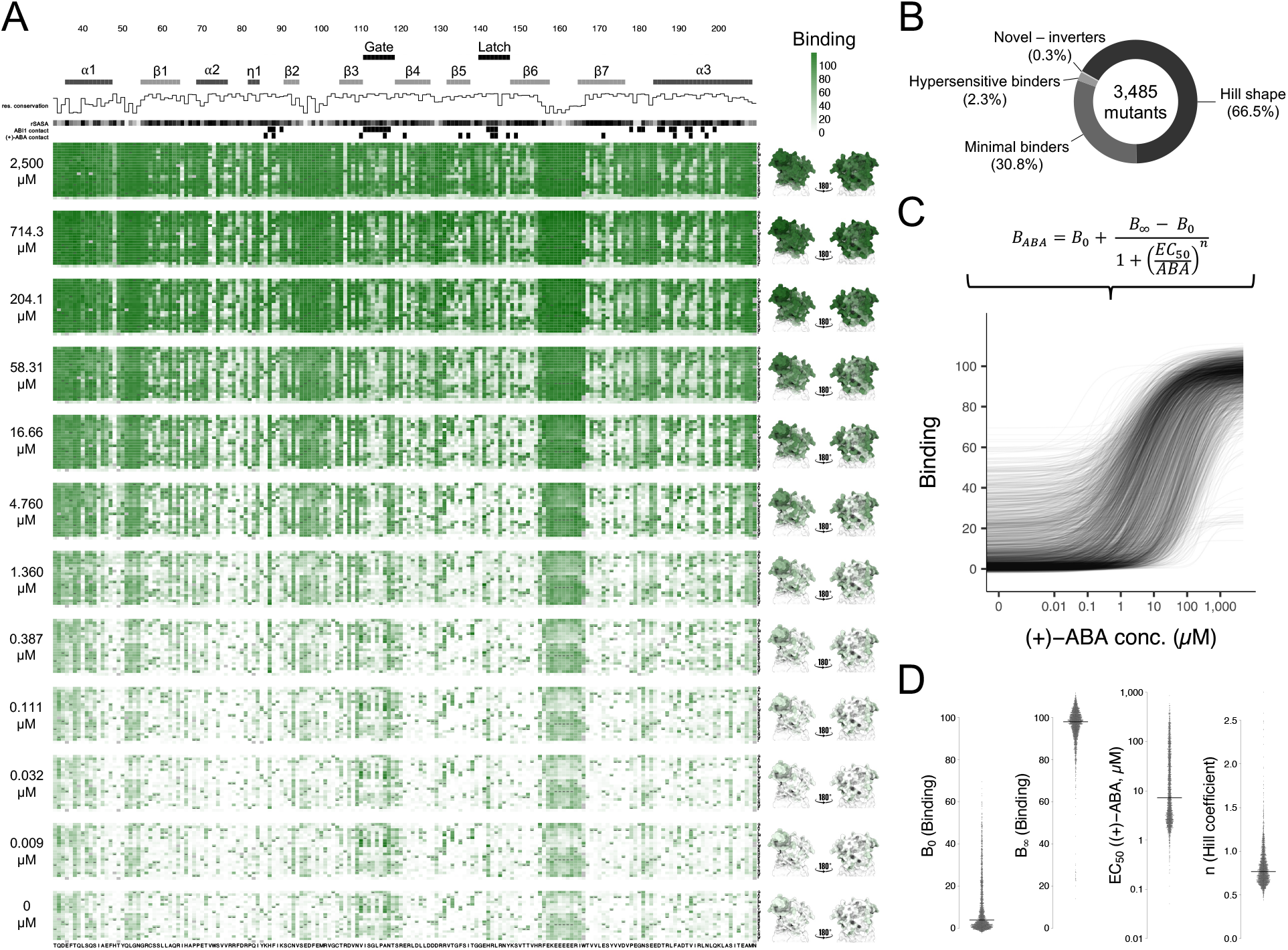
Hormone dose response profiles of thousands of PYL1 receptor mutants. **A:** ABI1-binding variant effect heatmaps of PYL1 mutants exposed to twelve (+)-ABA concentrations. Top layer: PYL1 sequence positions, secondary structure elements, residue conservation, residue solvent accessible surface area (SASA); residue positions in the ABI1 and hormone interfaces are annotated in accordance with Miyazono *et al*., 2009. SASA colour grey scaling is from lowest (black) to highest (light grey). Each heatmap features the PYL1 receptor amino acid sequence along the X-axis, with columns showing the PYL1-ABI1 binding fitness values of all 20 mutant genotypes per residue. Wildtype genotypes in each residue are depicted with a hyphen. The associated (+)-ABA concentration is labelled to the left of each heatmap, whereas mean residue-level variant binding is projected on the PYL1-ABI1 complex structure. **B:** Distribution of all PYL1 mutant dose response profiles with complete data, as determined by hierarchical clustering (see Supplementary Fig. S3A). **C:** Hill model fitting of PYL1 variant dose-response curves. Top: Hill equation as a function of (+)-ABA concentration. *B*_*0*_: binding at zero (+)-ABA concentration, *B*_*∞*_: binding at maximum (+)-ABA concentration, *EC*_*50*_: (+)-ABA concentration at 50% binding, *n*: Hill coefficient of curve steepness. Bottom: Display of 2,324 high-quality Hill curves (*R*^*2*^ > 0.95, *P*_*EC50*_ < 0.05, *P*_*n*_ < 0.05; all variant-level Hill curves can be found in Supplementary Fig. S4). **D:** Distributions of the inferred PYL1 mutant Hill parameters from high-quality dose-response curves (*R*^*2*^ > 0.95, *P*_*EC50*_ < 0.05, *P*_*n*_ < 0.05). Horizontal lines indicate median values.

### Mutations in 17.5% of residues increase receptor sensitivity (*EC*_*50*_)

Hierarchical clustering of the dose-response profiles for 3,485 PYL1 variants with complete measurements identified four major classes: 2,319 (66.5%) have sigmoidal dose-response curves; 1,075 (30.8%) show very little binding; 80 (2.3%) have hypersensitive binding, including constitutively in the absence of ligand; and 11 (0.3%) show novel inverted dose-response relationships (Fig. 2B, Supplementary Fig. S3A; Supplementary Table S3). Previously studied mutants behave as expected, including K86A (*41*) (minimal binding), H87P (*29, 33*), E141D (*44*) and A190V (*63*) (hypersensitive binding). To characterise the quantitative effects of each PYL1 mutation, we fitted Hill models to each variant’s dose-response profile, yielding 3,502 curves (Fig. 2C, dose-response curve for every mutation in Supplementary Fig. S4).

Distributions of all four Hill curve parameters vary substantially across genotypes (Fig. 2D): GluePCA *EC*_*50*_ values span over four orders of magnitude with an interquartile range (IQR) of = 2.81 µM to 39.74 µM (+)-ABA; *B*_*0*_ IQR = 1.00% to 15.22%; *B*_*∞*_ IQR = 93.47% to 101.31%; and *n* IQR = 0.67 to 0.90. Re-testing individual mutants validated the accuracy of the *EC*_*50*_, *B*_*0*_ and *B*_*∞*_ parameter fits (Supplementary Fig. S3B), and *EC*_*50*_ estimates are also concordant with an independent yeast-two-hybrid screen on the close homologue PYR1 (*63*) (Supplementary Fig. S3C).

Altogether, 91.4% of PYL1 mutants differ significantly from the receptor wildtype dose-response curve in at least one of the four Hill parameters (two-tailed *Z*-test, false discovery rate (FDR) < 0.01). Moreover, in 93.8% of residues three or more missense mutants have *EC*_*50*_ values that differ significantly from the PYL1 wildtype (two-tailed *Z*-test, FDR < 0.1). Most of these mutations increase *EC*_*50*_, yet dose responses of 79 variants in 31 positions (17.5% of residues) show marked gains in (+)-ABA sensitivity (Fig. 3A; Supplementary Table S3). In addition to known hormone interface or PPI residues, the comprehensive map reveals 22 novel sites where single mutations increase sensitivity, including multiple distal positions. One spatial cluster of these allosteric mutations includes substitutions at R164, I165 and W166 which lie >20Å from both the hormone pocket and PPI, in close proximity to the partially unstructured ß6-ß7 loop (Fig. 3B-E, Supplementary Fig. S3D).

**Figure 3:**
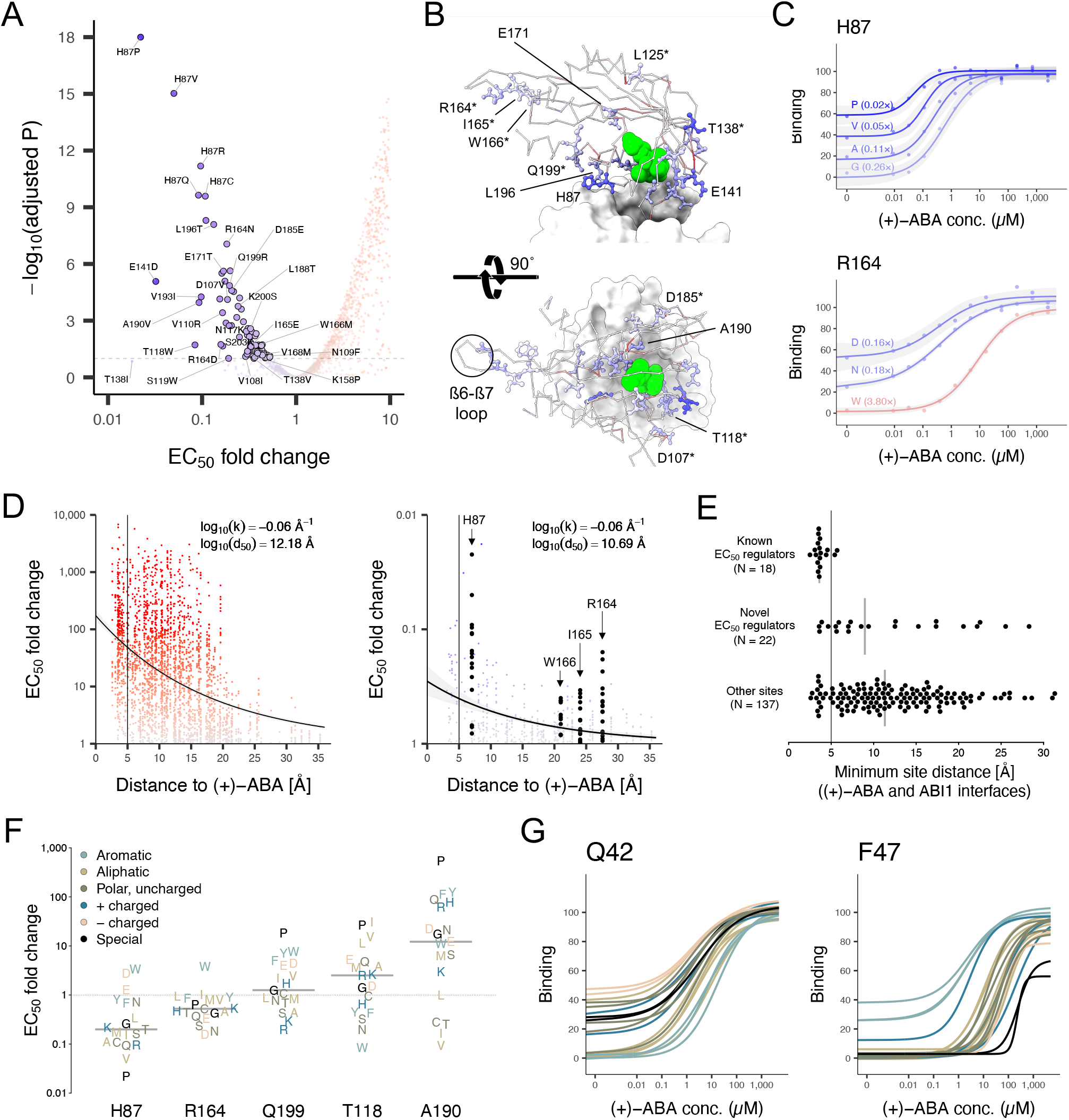
A complete allosteric mutational tuning map of PYL1 receptor sensitivity. **A:** Volcano plot, displaying inferred *EC*_*50*_ fold-changes between mutants and wildtype PYL1 on the X-axis against mutants’ inverse logarithmic *P* value (two-tailed *Z*-test) on the Y-axis, adjusted for multiple testing by false-discovery rate (FDR) control. Data point with *P* = 0 was set to 1×10^−18^. Colour scaling from blue to red is linear along the X-axis. Dashed horizontal line depicts FDR < 0.1. Variants with *EC*_*50*_ < 1 and FDR < 0.1 are highlighted as bold data points (N = 79). **B:** Side- and top-view of the PYL1-ABI1 structure, highlighting example side chains of PYL1 receptor positions at which hormone sensitivity can be significantly increased by mutations. Bound (+)-ABA molecule is depicted in green, ABI1 surface is displayed in light grey. Each residue is coloured by the mutation with strongest *EC*_*50*_ decrease (e.g. H87P in case of H87). Residues of interest are labelled, and novel tuning sites identified in this screen are highlighted with an asterisk. The ß6-ß7 loop, flanked by residues R164, I165 and W166, is encircled with a dotted line. **C:** Example dose-response curves of mutants in the sensitivity modulating interface residue H87 (top) and in the here described allosteric modulator residue R164 (bottom). Relative *EC*_*50*_ fold-changes to the wildtype are displayed above each mutant’s regression line. Grey shading indicates the 95% confidence interval of Hill model regressions. **D:** Euclidean distance decay scatterplot of *EC*_*50*_ fold changes within the PYL1 structure. X-axes indicate measured distances between receptor positions side-chain heavy atoms and the closest hormone atom. Y-axes indicate absolute *EC*_*50*_ fold changes, in order to directly compare the scales of increases (left panel with red-scaled data points) and decreases (right panel with blue-scaled data points). Bold black data points indicate particular sites of interest. Black curves depict regressions of inferred exponential decay, grey shadings indicate the 95% confidence intervals. Vertical black lines at X = 5 Å depict approximate contacts to (+)-ABA. Key rate parameter estimates are shown for both regressions. **E:** Summary of PYL1 sites in which single mutations can significantly decrease *EC*_*50*_. Minimum residue distances to (+)-ABA and ABI1 are depicted on the X-axis. The three data splits indicate known *EC*_*50*_-decreaser ‘regulator’ sites (N = 18), novel sites here described (N = 22) and all other sites (N = 137). Grey vertical bars indicate the mean minimal interface distances per set. **F:** Amino acid chemotype-resolved representation of selected PYL1 sites which can be altered towards either higher or lower hormone sensitivity by single mutations. Grey horizontal bars indicate mean *EC*_*50*_ fold changes per site. **G:** Example dose-response curve summaries of two PYL1 residues, in which mutants are coloured by their amino acid side chain chemotype as in (F) (see Supplementary Fig. S4).

### Distance-dependent decay of allosteric gain-of-function mutations

Receptor gain-of-function variants with decreased *EC*_*50*_ are enriched in the hormone pocket and ABI1-contacts, with a striking distance-dependent decay in effect with increasing Euclidean distance to the binding interfaces (Fig. 3D, Supplementary Fig. S3D). This widely-dispersed, yet approximately exponential distance-dependent decay is reminiscent of the distance-dependent decay of mutational effects on binding energy quantified in protein binding domains (*52, 53*) and also on the activity of the Src kinase (*54*). However, while the trend was only observed for loss- of-function variants in these examples, (+)-ABA distance-dependent decay is also seen for gain- of-function mutations in PYL1, with a log_10_(50%) reduction in mutational effects on *EC*_*50*_ at distances of 12.2 Å and 10.7 Å for loss- and gain-of-function, respectively (Fig. 3D).

Hill modelling of receptor mutants further shows that site-specific *EC*_*50*_ distributions are often governed by simple chemotype or side chain size preferences (Fig. 3F-G, Supplementary Fig. 4). For example, (+)-ABA sensitivity can be substantially increased via substitution of T118 to aromatic or polar, uncharged side chains. Altogether, these observations highlight a vast discovery potential for rational receptor tuning beyond alanine or glycine scanning.

### Correlated changes in sensitivity, maximal response, basal activity and steepness

Strikingly, across the PYL1 mutants, changes in the four Hill parameters are correlated (Fig. 4A). For example, *B*_*0*_ and *B*_*∞*_ are strongly – but non-linearly – correlated (Spearman’s *ρ* = 0.50), whereas *B*_*0*_ and *EC*_*50*_ (*ρ* = -0.63) and *B*_*∞*_ and *EC*_*50*_ (*ρ* = -0.66) are anti-correlated. These associations suggest the possibility of a shared underlying causal mechanism. Performing principal component analysis on the dose-response curves reveals that 73.9% of variation contained in the dose-response signals is captured by a single principal component (PC1; Fig. 4B), further suggesting that a common molecular trait drives most of the quantitative changes in Hill parameters.

**Figure 4:**
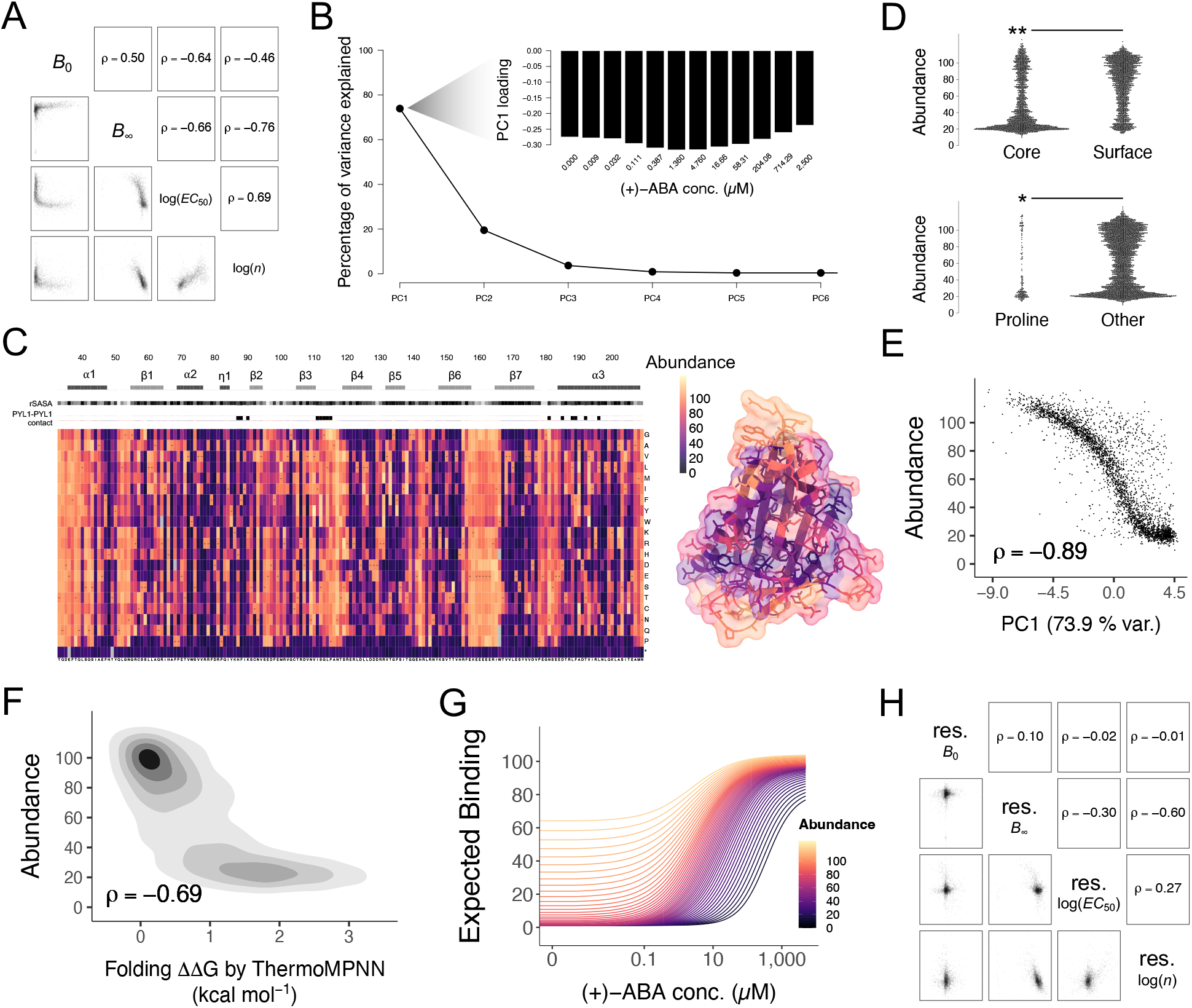
Receptor protein abundance is central to dose-response curve tuning. **A:** Pair-wise scatter plots and Spearman’s rank coefficients between Hill parameters inferred from high-quality dose-response curves (*R*^*2*^ > 0.95, *P*_*EC50*_ < 0.05, *P*_*n*_ < 0.05). *ρ*, Spearman’s rank coefficient. **B:** Scree plot of the explained variance by the first principal components (PC) that reduce the dimensionality of the complete set of PYL1 missense (N = 3,315) dose-responses. PC1 accounts for 73.9% of variance in the data. Inset bar plot of PC1 loading highlights an even contribution across (+)-ABA concentrations. **C:** Left: PYL1 abundance variant effect heatmap at 0 µM (+)-ABA (see Supplementary Fig. S5A). Top layer: secondary structure elements, relative SASA, structural positions in the PYL1-PYL1 interface are annotated. SASA colour grey scaling is from lowest (black) to 1 (light grey). Right: mean variant abundance per residue is projected on the Apo-PYL1 structure. **D:** Top: Distribution of mutant abundance effects in the PYL1 receptor core and surface. Bottom: Distribution of mutant abundance effects in PYL1 receptor proline sites and other amino acids. *one-sided Mann-Whitney U test *P* < 1×10^−11^, ** *P* < 1×10^−14^. **E:** Scatterplot of measured PYL1 mutant abundance versus the first principal component from dose-response curve dimensionality reduction (see B). **F:** Scatterplot of measured PYL1 mutant abundance versus PYL1 mutant ΔΔG_*f*_ predicted by ThermoMPNN (see C). *ρ*, Spearman’s rank coefficient. **G:** Theoretical dose-response curve distribution of PYL1-ABI1 binding upon variation of receptor abundance only. Model parametrisation was based on locally estimated scatterplot smoothing (LOESS) regression estimates between abundance and all four Hill parameters (see Materials and Methods). **H:** Pair-wise scatter plots and Spearman’s rank coefficients between residual distances of measured PYL1 abundance to the four Hill parameters inferred from high-quality dose-response curves (*R*^*2*^ > 0.95, *P*_*EC50*_ < 0.05, *P*_*n*_ < 0.05). Correcting for receptor abundance results in a strong decrease of correlations relative to those of the raw Hill parameters (see A) (see Supplementary Fig. 5B). *ρ*, Spearman’s rank coefficient.

### Receptor stability is the latent variable driving correlated phenotypic change

Recent large-scale testing has confirmed that high numbers of missense variants typically destabilize proteins to reduce their abundance (*50, 64*–*66*), including ∼60% of human pathogenic variants (*50*). Moreover, theoretical models predict that stability changes are one of several biophysical mutational effects that can drive correlated changes in dose-response curves (*8, 9, 67*). We hypothesized therefore that changes in receptor stability might be the underlying biophysical variable driving nearly 75% of the quantitative changes in Hill parameters.

To independently quantify the effects of mutations on the stability of PYL1, we used an orthogonal protein complementation assay that reports on the cellular abundance of PYL1 homodimers (*25, 29*) (Fig. 4C, Supplementary Fig. S5A). 2,690 missense variants (80.0%) reduce the abundance of mutant PYL1 - wildtype PYL1 dimers (one-tailed *Z*-test, FDR < 0.1), with mutations in the hydrophobic core and secondary structure elements particularly detrimental, as expected for a globular protein. Also as expected, proline mutations are detrimental throughout secondary structure elements, with alpha-helices and beta-sheets otherwise displaying their expected characteristic periodic patterns of mutational tolerance (*50, 64*) (Fig. 4C-D). These PYL1 homodimer abundance measurements were near-identical when cells were grown either in the total absence or presence of 250 µM (+)-ABA (*ρ* = 0.99; Supplementary Table S2).

Strikingly, the PYL1 mutant abundance measurements were very strongly correlated with the first principal component of the PYL1-ABI1 binding dose response measurements (*ρ* = -0.89; Fig. 4E) as were *in silico* predictions of stability changes (*68*) (Fig. 4F; Supplementary Table S4), strongly supporting our hypothesis that changes in stability underlie the correlated changes in receptor parameters. To further quantify this, we plotted the abundance of each genotype against the four Hill curve parameters from 2,324 high-quality dose response curves (Hill model *R*^*2*^ > 0.95, *P*_*EC50*_ < 0.05, *P*_*n*_ < 0.05). Changes in all four parameters are tightly coupled to abundance changes (Supplementary Fig. S5B). Destabilized PYL1 variants have reduced sensitivity (increased *EC*_*50*_, *ρ* = -0.77) and reduced maximum response (decreased *B*_*∞*_, *ρ* = 0.60). Stable PYL1 variants, on the other hand, have higher baseline activity (increased *B*_*0*_, *ρ* = 0.81) and flatter induction profiles (decreased n, *ρ* = -0.57) (Fig. 4G). Changes in receptor stability are therefore the major mechanism by which missense variants drive correlated changes in dose-response curve parameters.

### Independent tuning of receptor signalling phenotypes

We next investigated changes in the four Hill curve parameters not accounted for by changes in PYL1 abundance. For each mutant we quantified the residual between each of its Hill parameter values and those predicted by locally estimated scatterplot smoothing (LOESS) regression between the parameter and abundance across all genotypes (Fig. 5A; Supplementary Table S5).

**Figure 5:**
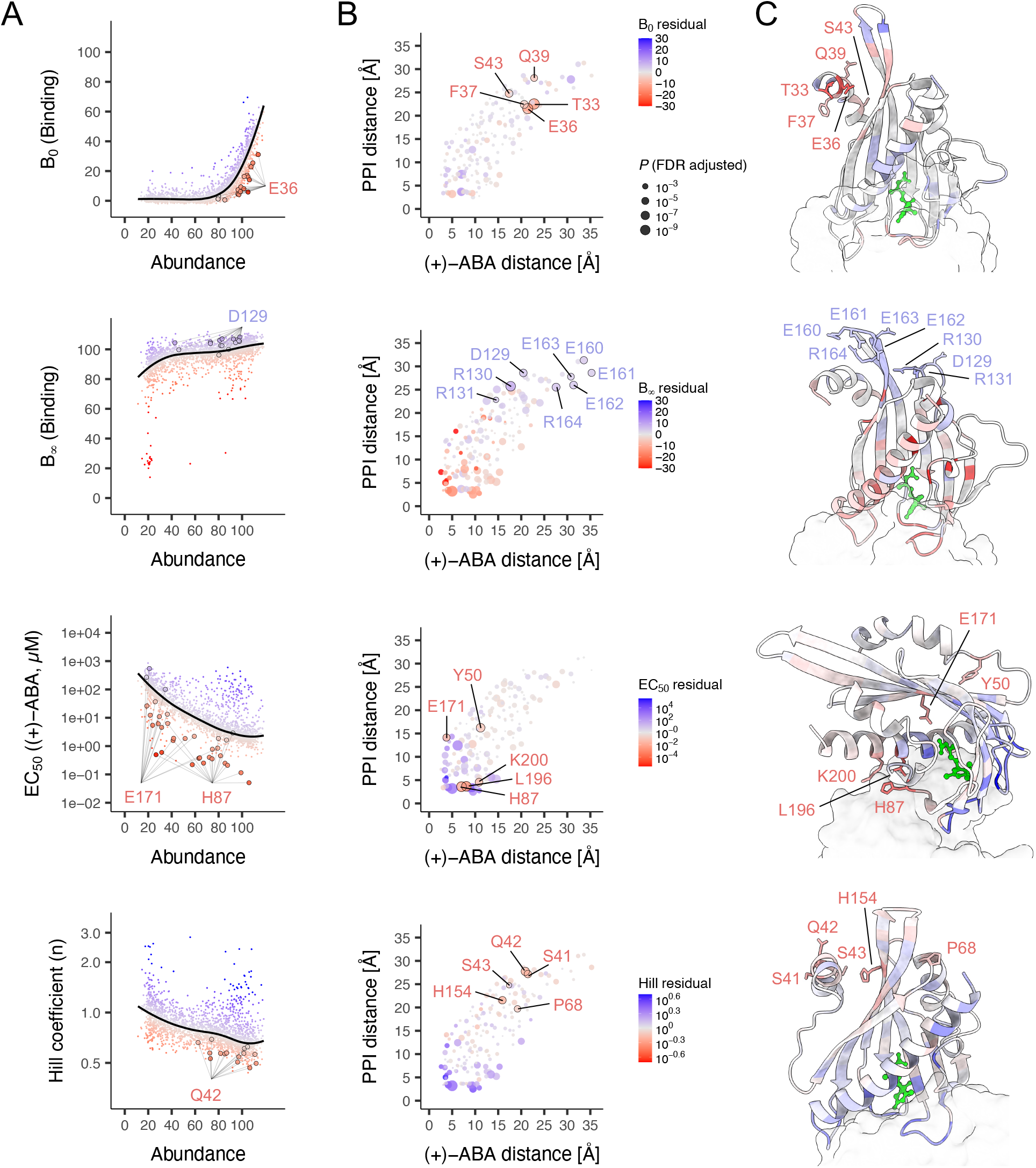
A highly resolved, modular landscape of receptor tuning after abundance normalisation. **A:** Scatterplots of mutant PYL1 abundance versus the four different Hill parameters of PYL1-ABI1 binding, with subpanels ordered from top to bottom: *B*_*0*_, *B*_*∞*_, *EC*_*50*_, *n*. Black lines in each subpanel indicate LOESS regressions, while data points represent individual mutants. Data points are colour scaled by their residual distances to the regression, from low (red) to high (blue). Encircled data points highlight mutations in example residues of PYL1 which are particularly amenable to Hill parameter tuning independent of protein abundance (see Supplementary Fig. S6A). **B:** Scatterplots of Euclidean distances of PYL1 residue side-chain heavy atoms to the nearest (+)-ABA (X-axes) and ABI1 (Y-axes) sites. Data points are coloured by their position-averaged mutant residual distance to Hill parameter LOESS regressions against abundance. Data point sizes represent -log_10_(*P*), thereb highlighting statistically significant positions as defined by two-sided Mann-Whitney U tests of pooled mutant residual distances to the regression of a particular PYL1 position vs. pooled mutant residual distances to the regression of all other positions, after multiple testing correction by false-discovery rate. Encircled data points highlight selected PYL1 positions with strong Hill parameter-unique tuning potential independent of protein abundance. Subpanel order and PYL1 colour scale as in (A) (see Supplementary Fig. S6B). **C:** Structural display of PYL1 mutant residual distance to Hill parameter LOESS regressions against abundance measurements. ABI1 surface is coloured in light grey. Subpanel order and PYL1 colour scale as in (A). Selected large- effect residues from (B) are labelled and side changes chains shown. Three-dimensional rotations of the complex were freely chosen to highlight relevant spatial clusters indicative of allostery Hill parameter regulation.

In total, mutations in 15, 44, 37 and 17 receptor residues had larger effects on *B*_*0*_, *B*_*∞*_, *EC*_*50*_ and *n*, respectively, than expected given their effects on abundance (Mann-Whitney U test, FDR < 0.01; Supplementary Table S5). Moreover, the mutational effects on *B*_*0*_, *B*_*∞*_ and *EC*_*50*_ are largely uncorrelated after accounting for abundance changes (Fig. 4H). Thus, while stability modulation drives correlated changes in the Hill parameters, mutational effects beyond stability allow for independent tuning of receptor behaviour.

### Modular structural encoding of receptor phenotypes

To better understand the mutations that independently tune each of the four Hill curve parameters, we visualised the mean abundance-corrected mutational effects on the atomic structure of PYL1 (Fig. 5B-C). This reveals a highly modular genetic architecture in which distinct regions of the protein fold can tune particular Hill curve parameters. Large-effect sites which can independently tune the basal and maximum receptor activity point towards two spatial clusters located in ligand- distal extensions of the PYL1 fold, either via the neighbouring ß4-ß5 and ß6-ß7 loops (*B*_*∞*_) or via the adjacent N-terminal helix (*B*_*0*_) (Fig. 5C).

Out of the six PYL1 positions where mutations cause the strongest decrease in basal binding beyond abundance changes, five (T33, E36, F37, Q39, S43) fall into the N-helix. In contrast, mutations that increase *B*_*∞*_ include multiple mutations in charged residues of the juxtaposed ß4- ß5 (D129, R130, R131) and ß6-ß7 (E160, E161, E162, E163, R164) loops. These two allosteric clusters are both located >15 Å from the key ternary complex interfaces, demonstrating the importance of long-range energetic interactions in receptor tuning (Fig. 5B, Supplementary Fig. S6A-B).

Closer to the hormone binding pocket, nearly all mutations in the (+)-ABA contacting residue E171 and the second-shell residue Y50 reduce *EC*_*50*_ more than expected after correcting for their effects on abundance (Fig. 5B, Supplementary Table S5). Interestingly, unlike the known *EC*_*50*_ regulator site H87, mutations in these two positions also strongly reduce receptor abundance (Fig. 5A, Supplementary Fig. S6A). Missense mutations in the (+)-ABA-E171-Y50 axis may thus point towards an evolutionarily trade-off between PYL1 stability and signalling.

### Phenotypic innovation

Finally, we examined the 91 receptor mutants (2.6%) with hypersensitive PYL1-ABI1 binding (Fig. 2B). Eleven of these have inverted dose-response curves in which ABI1 interactions are reduced at higher (+)-ABA concentration, thereby transforming the receptor from an OFF-ON into an ON-OFF switch (Fig. 6A). Re-testing individual mutants confirmed the inverted phenotypes (Fig. 6B), and with negligible effects on abundance (Fig. 6C).

**Figure 6:**
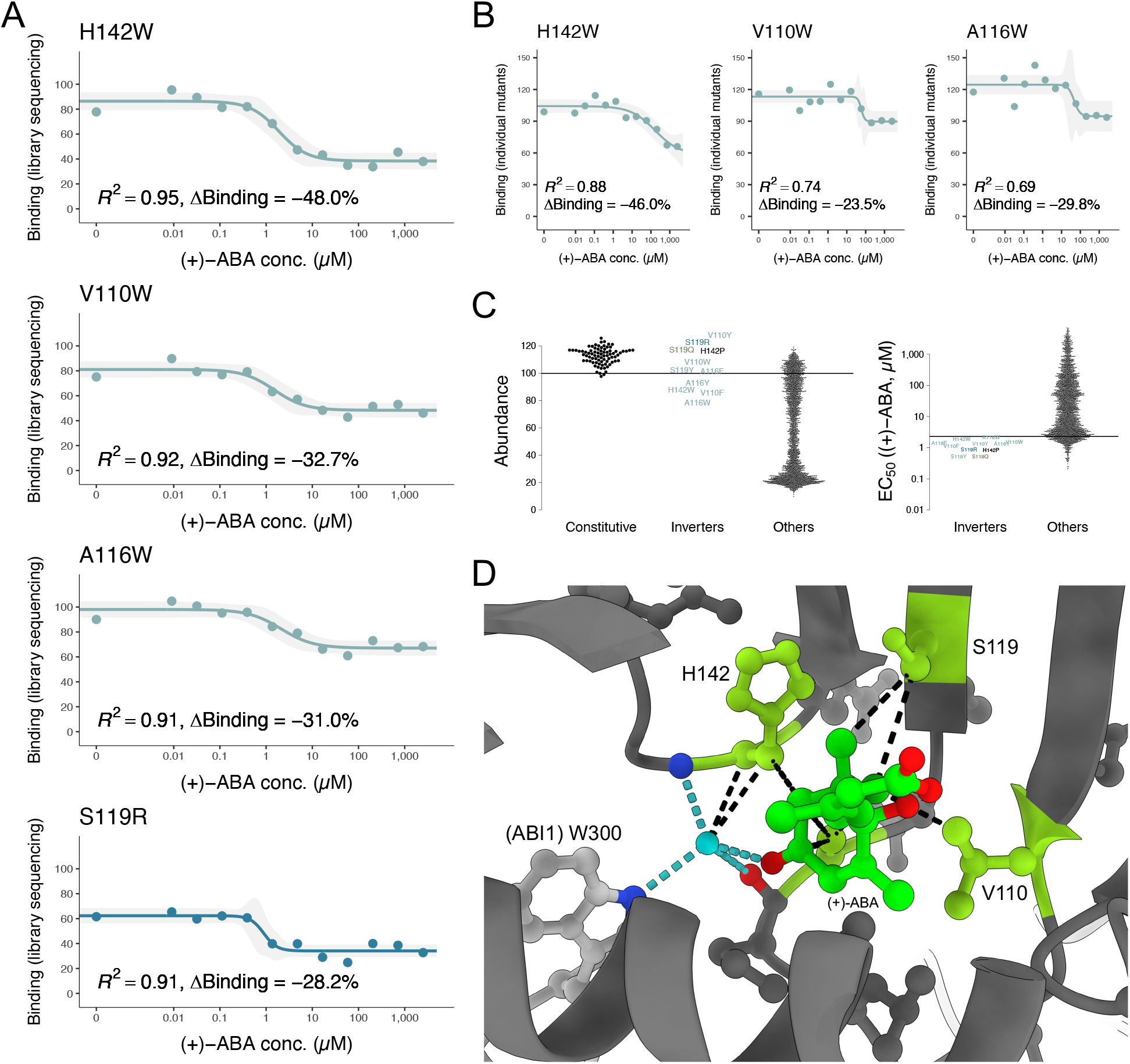
0.3% of PYL1 single mutants result in a novel inverted dose-response phenotype. **A:** Inverted dose-response curves of example mutants for which the OFF-switch phenotype was identified by bulk GluePCA. Grey shading represents 95% confidence interval. *R*^*2*^, coefficient of determination. ΔBinding, relative binding difference between estimated *B*_*0*_ and *B*_*∞*_. **B:** Manual validation of the three aromatic, dose-response inverting mutants with strongest binding decrease in the original deep mutational scanning library experiment. Grey shading represents 95% confidence interval. *R*^*2*^, coefficient of determination. ΔBinding, relative binding difference between estimated *B*_*0*_ and *B*_*∞*_. **C:** Left: distributions of mutant abundance in constitutive, ‘inverter’ and all other PYL1 receptor mutants. Horizontal line represents PYL1 wildtype abundance. Right: distributions of mutant *EC*_*50*_ in inverter and all other PYL1 receptor mutants. Horizontal line represents PYL1 wildtype *EC*_*50*_. **D:** Atomic-level structural view of the (+)-ABA binding pocket, in light green highlighting the three PYL1 dose-response inversion mutant featuring residues V110, S119 and H142. PYL1 secondary structure elements are represented in dark grey, ABI1 in light grey. Side chains of other receptor residues which feature constitutive binding mutants where observed are also displayed in dark grey, as is the ABI1 W300 ‘tryptophan lock’ residue in light grey. The W300 indole N-H (dark blue) lies at the coordinative base of a key water molecule (cyan), stabilised by additional Hydrogen bonds (cyan, dashed lines) with the backbone amide (dark blue) of PYL1 gate loop R143, the peptide carbonyl oxygen of latch loop P115 (red), and the C4 carbonyl oxygen of (+)-ABA (red). Van der Waals forces involving the PYL1 inverter sites are displayed as black dashed lines. In the here chosen representation, (+)-ABA (dark green) overlays A116, the fourth residue in which multiple inverted mutant phenotypes were observed.

All eleven PYL1 ‘inverter’ mutations coincide with gate (V110, A116, S119) and latch-associated (H142) loop positions (Fig. 1B) whose wildtype side chains form hydrophobic contacts with the chemical’s cyclohexene ring, its C6 methyl and C4 carbonyl groups (Fig. 6D); eight of these are substitutions to bulky aromatic amino acids (Fig. 6A-C). The latch loop variant H142W features the strongest relative binding decrease (*B*_*0*_ = 86.4, *B*_*∞*_ = 38.4; Fig. 6A), whereas the majority of inverter mutants display moderate effects (median ΔBinding = -26.9%).

We note that both A116 and H142 are located in direct vicinity to residues which coordinate the key water molecule orientation in gate-latch closure (*69*) (Fig. 6D). Aromatic or hydrophobic mutations at the equivalent (+)-ABA pocket positions of V108, V110, I111, S112, L114, A116, H142 and R143 have been previously reported to confer constitutive or hypersensitive binding in the close PYL1 homologue PYR1 (*33, 41, 44, 63*) but not, to our knowledge, inverted responses. Interestingly, the *EC*_*50*_ values of inverters are similar to those of the PYL1 wild type (Fig. 6C). This suggests a relatively simple mechanistic flip, potentially due to steric displacements within the ligand binding pocket that are further exacerbated upon hormone addition.

## Discussion

We have generated here a complete site-saturation map of how amino acid substitutions quantitatively and qualitatively alter the input-output signal processing of a receptor protein.

Our comprehensive mutagenesis and phenotyping of the PYL1 drought sensor generated complete dose-response curves for >3,500 different proteins. The data reveal that at least 91% of all amino acid substitutions in this hormone receptor quantitatively alter its input-output function, and lead to the identification of 22 novel receptor sites at which single mutations can increase hormone sensitivity – more than doubling the number reported thus far. Moreover, we show that ∼75% of the changes in signalling are caused by protein stability changes. Stability changes have not been accurately quantified in previous medium- and large-scale mutagenesis studies of receptors (*13*–*15*) and both our data and theoretical work (*8, 9, 67*) predict that many of the reported changes in dose-response curves in these previous studies will be caused by latent changes in receptor stability.

Strikingly similar to what has been reported for small protein-binding domains and the kinase Src, the allosteric effects of mutations on the *EC*_*50*_ of the PYL1-(+)-ABA-ABI1 interaction show a strong distance decay away from the interface. This suggests that distance-dependent allosteric decay is a conserved principle of protein biophysics. Uniquely for PYL1, this allosteric decay is observed for both loss-of-function and gain-of-function mutations. In future work it will be important to quantify allosteric communication in additional protein-small molecule complexes across a multitude of binding modes and interface geometries, to evaluate the generality of this finding and to better understand the principles of how mutations influence small molecule sensing by proteins.

Changes in stability drive correlated changes in multiple signalling parameters and are the most frequent biophysical effect of mutations, particularly in buried hydrophobic cores (*50, 55, 64*). In contrast, our data show that mutational effects on signalling beyond stability changes have a different genetic architecture, with mutations in different structural regions tuning *B*_*0*_, *B*_*∞*_ and *EC*_*50*_. The receptor’s genetic architecture is thus modular, allowing independent control of signalling parameters by sequence changes in different structural modules. We note that mutations in the partially unresolved, interface-distal ß6-ß7 loop of PYL1 have an unexpectedly strong influence on raw hormone sensitivity and stability-independent *B*_*∞*_. Moreover, several positions in the N-terminal helix serve as independent regulators of basal PYL1-ABI1 binding. We propose that the conformational dynamics of such flexible regions may yield entropic reservoirs which contribute to the free energy of CID complex formation (*70*), long-range mechanisms which warrant systematic enquiry across receptor-ligand pairs.

Beyond these quantitative changes, we were also surprised to identify individual missense variants with qualitative changes in signalling. These phenotypic innovations are rare, highlighting the discovery power of complete saturation mutagenesis, and illustrate how even single amino acid substitutions can dramatically alter the allosteric response of a receptor switch. In total, eleven variants in PYL1 invert its dose-response curve and transform the protein from an OFF-ON to an ON-OFF switch. Previous work has demonstrated the remarkable ability of mutations in the ligand pocket of PYL/PYR proteins to reprogram their specificity towards new ligands (*34*– *39*). PYL1 is thus a highly tunable and evolvable allosteric protein. Complete mutagenesis and quantitative phenotyping of additional protein families will be required to understand if this malleability is unusual or a common feature of receptors. The huge evolutionary expansion of other receptor families such as GPCRs (*71*) and nuclear receptors (*72*) suggests that the latter is likely to be the case, as does the diversity of chemical ligands that can specifically bind prokaryotic allosteric TFs and their quantitative responses to mutagenesis (*73*).

We believe that the approach that we have taken here can now be applied to many other chemically-inducible protein switches. The specific experimental method used in this work – GluePCA – may be applied to other soluble receptors and drug-mediated dimerization systems (Supplementary Fig. S1). Moreover, related pooled selection experiments have been developed for cell surface membrane receptors (*74*–*78*), although not yet applied at scale to generate full ligand dose-response curves. Combined with selections for biophysical properties, this approach should allow the deep exploration of protein genotype-phenotype encoding and a mechanistic understanding of how protein sequences encode signalling parameters.

## Acknowledgements

This work was funded by Wellcome (220540/Z/20/A), the European Research Council (ERC, Advanced Grant 883742), the Spanish Ministry of Science and Innovation (LCF/PR/HR21/52410004, EMBL Partnership, Severo Ochoa Centre of Excellence), AGAUR (2021 SGR 01226), and the CERCA Program/Generalitat de Catalunya. M.R.S. was funded by an EMBO Long-term Postdoctoral Fellowship (ALTF 544-2021) and by the European Union’s Horizon 2020 research and innovation programme under the Marie Skłodowska-Curie grant ‘DeepGlue’ (101107635).

We thank members of the Lehner laboratory, Rosa Martínez-Corral, Adrian Baez-Ortega, Pius Galli, Nicolas Thomä and Tim Whitehead. We thank Antoni Beltrán for ThermoMPNN calculations. The *TOR1-1* Δ*fpr1* yeast strain used for rapamycin GluePCA was a generous gift by Anuj Kumar (University of Michigan). We thank the CRG Genomics Unit and Wellcome Sanger Institute DNAP for DNA sequencing services.

## Data availability

All DNA sequencing data have been deposited in the European Nucleotide Archive (ENA) under project accession no. PRJEB89674.

## Code availability

Source code to reproduce the analyses is available at https://github.com/lehner-lab/ABA_receptor

## Author contributions

M.R.S. performed all experiments, analyses and wrote the original manuscript. M.R.S. and B.L. conceived experiments, analyses and wrote the final manuscript.

