## Supplementary Methods for "The genetic architecture of an allosteric hormone receptor"

### 1. Experimental methods

#### 1.1 Yeast inducible Protein Complementation Assay (GluePCA) vector design

Wildtype *Arabidopsis thaliana* PYL1 (residues 33-209 (*40*)) and ABI1 (residues 126-423 (*40*)) protein-coding DNA sequences were obtained through Addgene (plasmids #135988, #135985 (*79*)). ABI1 was amplified with primers oMS093_fw and oMS093_rv, introducing in-frame overhangs for enzymatic NheI and HindIII restriction sites; PYL1 was amplified with primers oMS094_fw and oMS094_rv, introducing in-frame overhangs for enzymatic BamHI and SpeI restriction sites (Supplementary Table S1A). Both amplicons were double-digested with their matched high-fidelity restriction enzymes (37°C, 5 hours) and subsequently cloned into the inducible Protein Complementation Assay (GluePCA) expression vector by T4 temperature-cycle ligation (*80*).

In this pMS-GluePCA1 plasmid, PYL1 and ABI1 are equally expressed in yeast cells by use of two separate, constitutive CYC1 promoters (Fig. 1C). PYL1 is transcribed as an N-terminal fusion to the N-terminal fragment of L23Y-F32S double mutant (*48*) murine, methotrexate (MTX) resistant dihydrofolate reductase (mDHFR residues 2-106 (*58*)). ABI1 is expressed as an N-terminal fusion to the C-terminal mDHFR fragment (residues 107-187 (*58*)). Both fusion proteins contain flexible $(GGGGS)_4$ linker sequences, and they share a common CYC1 transcriptional terminator (Supplementary Table S1B).

The pMS-GluePCA1 vector was designed so that (+)-ABA inducible *in vivo* protein-protein interactions between PYL1 and ABI1 also lead to the linked reconstitution of full-length mDHFR protein. Conceptually, this engineered mDHFR enzyme's function is essential for *Saccharomyces cerevisiae* survival in MTX-enriched medium since the cells' native scDHFR enzyme is inhibited in MTX. It follows that PYL1 mutants' differential potential to form a complex with ABI1 can be quantitatively inferred from their cellular growth rate mediated by functional mDHFR (*48*–*56*, *58*).

For the other tested yeast GluePCA plasmids, we replaced PYL1 with *Arabidopsis thaliana* GID1A (residues 2-345 (*57*); from an IDT gBlock) or the human FRB domain of mTOR (residues 2025-2114 (*58*); from Addgene plasmid #130812 (*81*)), and ABI1 with *Arabidopsis thaliana* GAI (residues 2-92 (*82*); from an IDT gBlock) or human FKBP12 (residues 2-108 (*58*); from Addgene plasmid #137831 (*81*)) (Supplementary Fig. S1; Supplementary Table S1B).

#### 1.2 Individual point mutagenesis

We used mutant forward primers or site-directed mutagenesis by overlap extension PCR (*83*) to engineer individual point mutants of PYL1 (Q34R, Q34I, E36R, H87A, H87V, H87P, A116S, T118N, T118W, S119A, S119V, S119N, H142C, L144A), ABI1 (W300A), and GAI (E54R) (Fig. 1E, Supplementary Fig. S1; Supplementary Table S1C).

The following point mutants were ordered as gBlocks sequences, amplified and cloned into their respective GluePCA vectors: PYL1 Q34Y, T38K, Q39F, H48C, D80M, P82G, Y85P, V110H, V110Y, V110W, A116H, A116W, A116R, T118I, L125I, V132P, I137G, H142W, E161A, E171K, A190E, V193H, V193W, R195M, A202T (Supplementary Fig. S3), as well as mTOR/FRB Y2088A-K2095P-T2098L-W2101F (*59*) (Supplementary Fig. S1).

#### 1.3 Small-scale chemically inducible dimerisation experiments

All GluePCA plasmids were chemically transformed into *Saccharomyces cerevisiae* strain BY4741 (PYL1-ABI1, GID1A-GAI) or the rapamycin-resistant *Saccharomyces cerevisiae* strain *TOR1-1 ∆fpr1* (*84*) (FKBP12-mTOR/FRB). Transformed cells were selected on agar plates containing synthetic complete (SC) yeast minimal medium without uracil (-URA) and adenine (-ADE). SC -URA -ADE plates were incubated at 30°C for 48 hours, to allow for the outgrowth of plasmid-carrying yeast colonies.

MTX (Sigma-Aldrich, St. Louis, USA; Catalog No. PHR1396) was dissolved in liquid Dimethyl Sulfoxide (Panreac Química SLU, Castellar de Vallès, Spain; Catalog No. 131954) to a final concentration of 10 mg/mL. This MTX stock solution was subsequently used to dissolve solid Abscisic Acid ((+)-ABA) (Selleck Chemicals LLC, Houston, USA; Catalog No. S7594), solid Gibberellic acid (GA3) (Selleck Chemicals LLC, Houston, USA; Catalog No. S4766) or solid Rapamycin (Selleck Chemicals LLC, Houston, USA; Catalog No. 1039) to stock concentrations of 125 mM, 125 mM and 50 mM, respectively. Subsequent dilutions of these small molecule inducer-enriched MTX stocks were done with the original 10 mg/mL MTX solution.

Two days after plasmid transformation, individual yeast colonies were inoculated into 150 µL of liquid SC -URA -ADE medium in sterile Corning 384 well deep-well plates (Sigma-Aldrich, St. Louis, USA; Catalog No. CLS3342). Cell suspensions were grown under constant shaking, at 30°C for 24 hours. After that, yeast cells were diluted 1:20 into fresh SC -URA -ADE medium. 4 µL of each dilution was then inoculated into 76 µL competition medium, composed of liquid SC -URA -ADE with 200 µg/mL MTX (*51*) and enriched with different concentrations of (+)-ABA, GA3 or rapamycin. These cultures were prepared within sterile, transparent, non-treated Nunclon 384 well flat-bottom microplates (Thermo Fisher Scientific Inc, Waltham, USA; Catalog No. 242765). Plates were tape-sealed and then transferred to an Infinite M200 PRO microplate reader (TECAN Group Ltd, Männedorf, Switzerland). Yeast cultures were grown at 30°C for 60-72 hours. Optical densities at 600 nm wavelength ($OD_{600}$) of each individual well was measured in 15 minute intervals, respectively followed by short intervals of vigorous shaking.

#### 1.4 PYL1-PYL1 homodimer abundancePCA vector design

To quantify PYL1-PYL1 homodimer interactions, we generated a derivative vector from pMS-GluePCA1 (Supplementary Table S1B).

We codon-optimised a second wildtype copy of PYL1 (residues 33-209 (*40*)) in two steps: (1) the overall nucleotide sequence was biased towards high translation efficiency in *Saccharomyces cerevisiae*, (2) nucleotide sequences of amino acids overlapping with any primer binding sites for amplification of the four PYL1 mutagenesis pools (see 1.5) were manually optimised to increase Hamming distance to the PYL1 in GluePCA. While identical on the amino acid level, the two PYL1 nucleotide sequences differ by 43.5%.

We digested, dephosphorylated and gel-purified the pMS-GluePCA1 vector with restriction enzymes NheI-HF, HindIII-HF and Quick calf intestinal alkaline phosphatase (37°C, 5 hours) (New England Biolabs, Ipswich, USA; Catalog No. R3131, R3104, M0525). The second, nucleotide-sequence recoded copy of wildtype PYL1 was then amplified from an IDT gBlock (Integrated DNA Technologies Inc, Coralville, USA) via polymerase chain reaction, using primers oMS279_fw and oMS279_rv (Supplementary Table S1A). We then used T4 temperature-cycle ligation (*80*) to insert the recoded PYL1.

#### 1.5 PYL1 site-saturation mutagenesis library cloning

Cassette site-saturation mutagenesis of PYL1 was performed as follows. Four groups of mutant PYL1 sequences were synthesised as single-stranded oligonucleotide pools (Integrated DNA Technologies Inc, Coralville, USA), each spanning blocks of 39-46 amino acid positions by tiled sets of degenerate NNK codons (Supplementary Table S1D). Site-saturation is reached since NNK represents 32 nucleotide combinations which encode for all 20 amino acids and also the TAG ‘amber’ stop codon.

Using complementing 3'-oligonucleotides, the four variant libraries were double-stranded by a single, long PCR cycle (94°C for 3 min., 50°C for 30 sec., 72°C for 5 min.), prior to purification and concentration by MinElute columns (Qiagen, Hilden, Germany; Catalog No. 28004). We separately cloned these into a linearised, PYL1 containing minimal vector by two-fragment Gibson assembly. All four PYL1 mutant libraries were electrically transformed into high-efficiency, 10-beta electrocompetent *E. coli* cells (New England Biolabs, Ipswich, USA; Catalog No. C3020), recovered at 37°C in 2 mL super optimal broth with catabolite repression (S.O.C.) medium for 1 hour and subsequently selected in 50 mL Luria-Bertani (LB) medium containing 50 µg/mL spectinomycin at 37°C. 16-18 hours later, cells were harvested by centrifugation at 8,000 g. We then purified the PYL1 library plasmid DNA using the Plasmid *Plus* Midi Kit (New England Biolabs, Ipswich, USA; Catalog No. 12945).

Library cloning and transformation efficiencies were estimated by plating 1:2,000 and 1:20,000 dilutions of the S.O.C.-recovered *E. coli* cell suspension onto agar plates with LB containing 50 µg/mL spectinomycin. Between 1.5 and 4.4 Million colony-forming units were obtained, representing mean programmed variant coverages of >1,500-fold.

As an orthogonal quality control of mutant library integration, shallow whole plasmid long-read sequencing was performed by Plasmidsaurus using Oxford Nanopore Technology. Raw sequencing reads were mapped against the PYL1 wildtype reference using minimap2 v2.26-r1175 (*85*). Alignments were visually examined using the integrated genomics viewer v2.12.3 (*86*) with high-quality single, double and triple-nucleotide mutant calls highlighted by setting the base quality shading scale boundaries to Phred scores of Q15 and Q25.

We subcloned the four mutant PYL1 libraries into both the pMS-GluePCA1 and pMS-PYL1-PYL1 homodimer abundancePCA plasmid by BamHI-HF and SpeI-HF (New England Biolabs, Ipswich, USA; Catalog No. R3136, R3133) digestion and gel purification, followed by overnight T4 temperature-cycle ligation (*80*). Libraries were electrically transformed and S.O.C. recovered as described before, and selection was performed in 50 mL LB medium containing 100 µg/mL ampicillin. Library cloning efficiencies and wildtype PYL1 vector religation carryover were estimated by plating of 1:2,000 and 1:20,000 dilutions onto agar plates with LB containing 100 µg/mL ampicillin. Mean variant coverages ranged at >50-fold, and consistently showed <1% wildtype plasmid carryover.

### 1.6 Bulk PYL1 mutant GluePCA competition across a 12-step, logarithmic (+)-ABA titration series

#### 1.6.1. PYL1 library transformation

For each of the four libraries, three individual BY4741 yeast colonies were inoculated into Erlenmeyer flasks with 20 mL yeast peptone dextrose adenine (YPDA) medium and shaken at 220 rounds per minute, at 30°C for 16 hours. To enter exponential growth, yeast cultures were then diluted to $OD_{600}$ of 0.3 in 100 mL fresh YPDA. After an additional 4 hours and reaching $OD_{600}$ of ~1.2, cells were prepared for chemical transformation by use of the high-efficiency lithium acetate method with single-stranded, sonicated Salmon sperm (Agilent Technologies Inc, Santa Clara, USA; Catalo No. 201190) carrier DNA and polyethylene glycol (LiAc/ssDNA/PEG) (*87*). For each transformation, 2 µg of pMS-GluePCA1 plasmid library DNA was added to the LiAc/ssDNA/PEG-buffered cells, prior to a 20 minute heat shock at 42°C. After an additional 1 hour of recovery, cells were transferred to 100 mL of fresh SC -URA -ADE medium. Erlenmeyer flasks with liquid yeast cultures at 30°C were shaken at 220 rounds per minute. After 19 hours, to enrich for live transformed cells the cultures were diluted to $OD_{600}$ of 0.2 in 100 mL of SC -URA -ADE medium ('input culture').

Library yeast transformation efficiencies were estimated by plating 1:1,000 dilutions of recovered cells onto SC -URA -ADE plates. Counts ranged consistently at >100,000 cfu, equivalent to mean PYL1 mutant coverages of >100-fold.

##### 1.6.2. (+)-ABA competition media preparation

We dissolved 125 mg of solid (+)-ABA (Selleck Chemicals LLC, Houston, USA; Catalog No. S7594) in 3,783 µL of DMSO previously enriched with 10 mg/mL MTX, yielding a 125 mM (+)-ABA stock solution as previously described (see 1.3). This (+)-ABA stock solution was manually diluted in ten steps, iteratively using the original DMSO stock solution with 10 mg/mL MTX at volumetric 1:2.5 ratios.

For each of the eleven (+)-ABA dilutions and an additional zero (+)-ABA control with only 10 mg/mL MTX, 1,632 µL (2% total volume) were dissolved in 80 mL of fresh SC -URA -ADE medium. The final (+)-ABA working concentrations were thus: 2,500.000 µM, 714.286 µM, 204.082 µM, 58.309 µM, 16.660 µM, 4.760 µM, 1.360 µM, 0.389 µM, 0.111 µM, 0.032 µM, 0.009 µM and 0.000 µM.

For all four PYL1 variant library pools to be tested against the exact same (+)-ABA concentrations, each competition medium was split into ~4 x 20 mL, kept in 100 mL Erlenmeyer flasks and pre-warmed to 30°C.

##### 1.6.3. PYL1 library GluePCA selection

After reaching the stationary growth phase ($OD_{600}$ ≥3.2), input cultures were inoculated at $OD_{600}$ of 0.05 into 20 mL competition medium (‘output culture’). Specifically, for each library the three input culture replicates were split so that they would be channeled into four different, stacked (+)-ABA concentrations across the dose range – replicate #1: 2,500.000 µM, 58.309 µM, 1.360 µM, 0.032 µM; replicate #2: 714.286 µM, 16.660 µM, 0.389 µM, 0.009 µM; replicate #3: 204.082 µM, 4.760 µM, 0.111 µM, 0.000 µM – and thereby yield a total set of twelve output cultures per variant library.

Remainder input culture cells were split into 50 mL Falcon tubes and spun down at 3,220 g for 5 minutes. Yeast cell pellets were twice washed with water, prior to being frozen at -20°C. Erlenmeyer flasks with output cultures at 30°C were continuously shaken at 220 rounds per minute. Depending on their bulk culture growth rate, after final $OD_{600}$ measurements cells were harvested 18.0 to 37.5 hours after inoculation. Similar to the input cultures, output cell pellets were washed twice with water and then stored at -20°C.

#### 1.7 Bulk PYL1-PYL1 homodimer abundancePCA competition

We transformed the four PYL1-PYL1 homodimer abundancePCA plasmid libraries into BY4741 yeast isolates and enriched for living cells, as described above (*87*) (see 1.6).

Two types of competition media were used for this selection experiment: SC -URA -ADE *(1)* with 2% DMSO containing 10 mg/mL MTX only, or *(2)* with 2% DMSO containing 10 mg/mL MTX and

250 µM (+)-ABA. Both sets of competition experiments were done in triplicates. Output cultures were grown for 25 hours prior to harvesting at final $OD_{600}$ of 1.4-2.0, equivalent to a mean of 4.8-5.3 yeast cell divisions in bulk culture inoculation.

### 1.8 Plasmid library DNA extraction

Plasmid library DNA extractions were performed using a custom phenol-chloroform protocol (*51*). Briefly, frozen yeast cell pellets from both input and output cultures were thawed at room temperature. We then resuspended the cells in 1 mL or 0.5 mL lysis buffer composed of 100 mM NaCl with 10 mM Tris-HCl, 1 mM EDTA, 2% Triton-X and 1% SDS for cell membrane disintegration.

In two cycles, cells were heated in a water bath at approximately 62°C for 10 minutes and then frozen in an ethanol bath with dry-ice at approximately -35°C for 10 minutes. We then added 1 g or 0.5 g of acid-washed 425–600 µm glass beads (Sigma-Aldrich, St. Louis, USA; Catalog No. G8772) and 1 mL or 0.5 mL of 25:24:1 phenol-chloroform-isoamyl alcohol (Sigma-Aldrich, St. Louis, USA; Catalog No. 77617). These suspensions were rigorously vortexed for 10 minutes at room temperature, prior to centrifugation at 3,220 g for 30 minutes. Aqueous upper phases of the centrifugates, containing library plasmid DNA fractions, were transferred to new Falcon tubes and complemented with the same amount of phenol-chloroform-isoamyl alcohol. After an additional 2-minute vortexing step at room temperature, the DNA suspensions were centrifuged at 3,220 g for 45 minutes. Aqueous phases of these secondary centrifugates were transferred to new Falcon tubes.

DNA precipitation was performed by complementation with 100 µL or 50 µL of 3M sodium acetate and 1.1 mL or 2.2 mL ice-cold molecular biology grade 100% ethanol (Panreac Química SLU, Castellar de Vallès, Spain; Catalog No. A8075). Precipitates were vigorously shaken and then incubated at -20°C overnight. To separate remaining phenol-chloroform traces and ethanol from the DNA, we centrifuged the isolates at 3,220 g and 4°C for 30 minutes.

Supernatant was carefully removed, and pellets air-dried at room temperature. DNA pellets were subsequently resuspended in 300 µL of Tris-EDTA buffer solution. We added 2.5 µL of RNAse A (Thermo Fisher Scientific Inc, Waltham, USA; Catalog No. K0502) and incubated the samples in a water bath at 37°C for one hour, prior to washing and bead-based DNA purification steps using the QIAEX II gel extraction kit (Qiagen, Hilden, Germany; Catalog No. 20051).

### 1.9 Plasmid DNA quantification and amplicon library preparation

We used quantitative PCR (qPCR) to estimate the number of library plasmid molecules in each yeast DNA isolate, using LightCycler SYBR green I master mix (Roche, Basel, Switzerland; Catalog No. 04707516001) and *ori*-binding primer sequences GCCTACATACCTCGCTCTGC and CAACCCGGTAAGACACGACT with 45 cycles of 95°C - 10 sec., 60°C - sec., 72°C - 10 sec (*51*). Dilutions of accurately pre-quantified wildtype GluePCA or PYL1-PYL1 plasmid were used to calibrate absolute estimates of library plasmid occurrences across experimental extracts. Based on the resulting qPCR values, we diluted the experimental input and output culture library DNA isolates in water to reach an even plasmid copy number across all samples.

Library PCR reactions were then set up to *(1)* specifically amplify only the mutated region of each isolate, *(2)* add 1-3 random 'N' nucleotide frameshifts to both the 5' and 3' end to increase library sequencing complexity, and *(3)* to attach generic Illumina i5 and i7 dual-index sequencing adaptor binding sites (*52*). Approximately $2.25 \times 10^7$ plasmid molecules, a mean >1,000-fold coverage per mutant, were used as inputs for each PCR reaction with 25 cycles of 94°C - 30 sec., 63°C - 30 sec., 72°C - 20 sec. using Q5 High-Fidelity Polymerase (New England Biolabs, Ipswich, USA; Catalog No. M0491). In order to remove excess oligonucleotides, PCR reactions were then treated with ExoSAP-IT enzyme (Thermo Fisher Scientific Inc, Waltham, USA; Catalog No. 75001) and incubated at 37°C for 1 hour, followed by a 20-minute heat inactivation at 80°C. Amplicons were extracted via QIAquick PCR purification columns (Qiagen, Hilden, Germany; Catalog No. 28104).

By use of a second PCR reaction, unique pairs of barcoded i5 and i7 Illumina PE sequencing adapters were ligated to each amplicon pool. We used 5 cycles of 94°C - 30 sec., 62°C - 30 sec., 72°C - 30 sec. using Q5 High-Fidelity Polymerase (New England Biolabs, Ipswich, USA; Catalog No. M0491). Resulting amplicons were subsequently purified using MinElute columns (Qiagen, Hilden, Germany; Catalog No. 28004).

#### 1.10 Next-generation sequencing

DNA concentrations from all barcoded amplicon libraries were measured using a NanoDrop One spectrophotometer (Thermo Fisher Scientific Inc, Waltham, USA; Catalog No. ND-ONE-W). Sequences from all samples were then diluted and pooled in an approximately equimolar ratio.

Pooled amplicons were deep-sequenced on an SP flow cell of a NovaSeq 6000 instrument, using paired-end sequencing at read length 150 bp (Illumina Inc, San Diego, USA; Catalog No. 20028400). An additional 10% PhiX DNA was spiked in to increase cluster complexity. Sequencing reads were subsequently de-multiplexed by barcode combination search using cutadapt.

Raw sequencing data for all samples has been deposited under the European Nucleotide Archive (ENA) project accession no. PRJEB89674.

### 2. Computational analyses

#### 2.1 Raw data processing and quality control

Raw read quality summarisation and filtering, sequencing adaptor and constant region trimming, alignment to the PYL1 reference sequence and mutant quantifications were performed using DiMSum v1.3.2 (*88*) (https://github.com/lehner-lab/DiMSum) with flags --vsearchMinQual 30, --fitnessMinInputCountAny 1:100, --fitnessMinInputCountAll 10, --mutagenesisType codon, --indels none, --maxSubstitutions 2, --mixedSubstitutions F. Using the --barcodeIdentityPath option, we moreover filtered for PYL1 variants with only exact matches to programmed NNK codons. To quantify the relative PYL1-ABI1 binding fitness of each PYL1 mutant *i* at each abscisic acid concentration, individual DiMSum calculations were performed for each of the 48 unique input-output culture combinations in GluePCA. PYL1-PYL1 homodimer fitness and errors were modelled using triplicate input-output matching.

Using these measures, across the full (+)-ABA dose response range we recovered PYL1-ABI1 binding fitness values for 3,511-3,529 out of 3,540 designed PYL1 amino acid variants in GluePCA sequencing data (99.2-99.7% programmed). From PYL1-PYL homodimer abundance measurements, we recovered fitness and error estimates for 3,533 out of 3,540 designed amino acid mutants (99.8%).

The DiMSum fitness $F_i$ of a mutant *i* is calculated as the natural logarithm of its sequencing counts from output culture $N_i^{output}$ over input culture $N_i^{input}$, and relative to the wildtype *wt* count (*88*):

$$F_i = \log\left(\frac{N_i^{output}}{N_i^{input}} \times \frac{N_{wt}^{output}}{N_{wt}^{input}}\right)$$

By design, such mutant fitness estimates are scaled relatively to the wildtype growth rate in each experimental condition. For improved comparability of GluePCA fitness values across the full twelve-step (+)-ABA titration, we therefore normalised all variants' relative sequencing counts to absolute per-hour growth rate estimates $GR_i$ by also integrating their associated experiments' recorded endpoint $OD_{600}$ measurement ($OD_{final}$) and total selection time *t*:

$$GR_i = \frac{\log\left(\dfrac{\dfrac{N_i^{output}}{\sum_i N^{output}} \times \mathrm{OD}_{final}}{\dfrac{N_i^{input}}{\sum_i N^{input}} \times 0.05}\right)}{t}$$

#### 2.2 Growth rate normalisation between library pools

By integration of growth rates from PYL1 stop ($GR_{stop}$) and synonymous wildtype ($GR_{syn}$) variants, which are expected to behave consistently between the four site-saturation mutagenesis libraries within each experimental condition, we calculated normalised mutant growth rates (GR)ˆ_i).

We first calculated the error ($\varepsilon$) weighted means $\mu_{GR_{stop}}$ and $\mu_{GR_{syn}}$, respectively:

$$\mu_{GR_{stop}} = \frac{\sum_{stop\, \in\, i} \frac{GR_{stop}}{(\varepsilon_{stop})^2}}{\sum_{stop\, \in\, i} \frac{1}{(\varepsilon_{stop})^2}} \qquad \mu_{GR_{syn}} = \frac{\sum_{syn\, \in\, i} \frac{GR_{syn}}{(\varepsilon_{syn})^2}}{\sum_{syn\, \in\, i} \frac{1}{(\varepsilon_{syn})^2}}$$

A linear affine transformation was then used so that $\mu_{GR_{stop}}$ and $\mu_{GR_{syn}}$ coincide between all four deep mutational scanning blocks, in all cases reference-calibrated with the first library pool's bimodal growth rate distribution centred on $\mu_{GR_{stop}}^{(1)}$ and $\mu_{GR_{syn}}^{(1)}$:

$$\widehat{GR}_i = \left(\frac{GR_i - \mu_{GR_{stop}}}{\mu_{GR_{syn}} - \mu_{GR_{stop}}}\right) \times \left(\mu_{GR_{syn}}^{(1)} - \mu_{GR_{stop}}^{(1)}\right) + \mu_{GR_{stop}}^{(1)}$$

#### 2.3 PYL1 mutant-specific relative binding and abundance scoring

Using the normalised iPCA growth rates of wildtype PYL1 across the full (+)-ABA dilution series ($\widehat{GR}_{WT}(ABA)$), we calculated a four-parametric Hill dose response curve by nonlinear least squares fitting with the *R* (*89*) drc package v3.0.1 (*90*), using the drm() function with presets fct = LL.4() and type = 'continuous' (see Fig. 1D):

$$\widehat{GR}_{WT}(ABA) = \widehat{GR}_{WT}(0) + \frac{\widehat{GR}_{WT}(\infty) - \widehat{GR}_{WT}(0)}{1 + \left(\frac{EC_{50}}{ABA}\right)^n}$$

The estimated maximum wildtype growth rate parameter ($\widehat{GR}_{WT}(\infty)$) was then used to rescale all other variants' normalised iPCA growth rates to relative $Binding_i$ scores:

$$Binding_i = 100 \times \frac{\widehat{GR}_i}{\widehat{GR}_{WT}(\infty)}$$

The PYL1 wildtype displays a residual ABI1 binding signal in the absence of (+)-ABA, an observation consistent between >200 independently tested synonymous wildtype genotypes in GluePCA (Fig. 1D, Supplementary Fig. S2A). High consistency between the relative binding of nonsynonymous PYL1 mutants in GluePCA and independent *in vitro* pull-down (*24*) (Fig. 1F) or yeast-two-hybrid (*63*) measurements (Supplementary Fig. S3C) suggests that the here chosen experimental design may be highly sensitised towards the detection of weak protein-protein interactions, as in the case of the PYL1 wildtype with ABI1. This may be due to the use of equal promoters (pCYC1) for the receptor and target protein components in GluePCA, while we note that *in vitro* ABI1 inhibition assays have been historically performed with up to 15-fold excess of receptor protein (*24*).

To also calculate $Abundance_i$ scores, we rescaled directly against the normalised WT growth rate:

$$Abundance_i = 100 \times \frac{\widehat{GR}_i}{\widehat{GR}_{WT}}$$

Supplementary table S2 contains all PYL1-ABI1 binding and PYL1-PYL1 abundance scores. Metadata and code to reproduce the calculations have been deposited in our Github repository: https://github.com/lehner-lab/ABA_receptor

#### 2.4 Dose response analyses

For classifications and hierarchical clustering of PYL1-ABI1 Binding phenotypes, we used 3,485 mutants – equivalent to 98.4% of programmed codons – for which fitness scores could be calculated across all twelve (+)-ABA concentrations tested (Fig. 2B, Supplementary Fig. S3A). We note that, while hierarchical clustering initially identified seven dose-response inverting mutants (V110F, V110Y, V110W, A116F, A116Y, A116W, H142W), further manual inspection of hypersensitive binders led us to switch the label of four additional mutants (S119R, S119Q, S119Y, H142P) to inverters.

An unbiased principal component analysis was performed by further removing 170 nonsense mutations (Fig. 4B). Specifically, we used the *R* prcomp()[87] function with value presets center = TRUE and scale. = TRUE.

PYL1 $Binding_i$ scores of all variants with complete data recovery across the twelve (+)-ABA dose response titration series were used to fit four-parametric log-logistic functions $f_{Binding_i}(ABA)$. Again, we used the *R* drc package v3.0.1 (*90*) with Hill parameters. $Binding_i(0)$, $Binding_i(\infty)$, $EC_{50_i}$ and $n_i$ were initialized on the wildtype PYL1 values. 95% confidence intervals of dose response curves were calculated using the default predict() function implemented in *R* drc v3.0.1 (*90*) across 1,000 evenly logarithmically spaced data points between 0 µM and 5,000 µM (+)-ABA. Nonlinear least squares ($R_i^2$) estimates of dose response curves were calculated by use of:

$$R_i^2 = 1 - \frac{\sum_{ABA}\left(Binding_i(ABA) - f_{Binding_i}(ABA)\right)^2}{\sum_{ABA}\left(Binding_i(ABA) - \overline{Binding_i}\right)^2}$$

Visual inspections led us to notice sets of mutant dose-response curves with unrealistically high $EC_{50}$ estimates, chiefly caused by their increased binding signal in only the highest one or two (+)-ABA concentrations. Other variants displayed constitutively low or high binding signals without a significant sigmoidal change across the dose-response titration series, resulting in flat Hill curve regressions with noisy $EC_{50}$ estimates. A third category of constitutively binding PYL1 mutants followed a non-monotonic dose-response, manifesting as band-stop phenotypes with lowest GluePCA signals at intermediate hormone concentrations (see the rightmost cluster in Supplementary Fig. S3A).

Thus, for in-depth Hill parameter comparisons and structural protein inferences, dose-response curves were quality-filtered by a number of conservative criteria: first, we removed dose response regressions with $R_i^2 < 0.95$. Secondly, we removed Hill models high $EC_{50}$ and $n$ parameter uncertainty ($P < 0.05$). This retained 2,337 dose response curves.

### 2.5 PYL1 binding vs. abundance residual analyses

For the high-quality, prefiltered dose response curves (see 2.4), we used locally estimated scatterplot smoothing regression to study the relationship between PYL1 abundance and each of the four Hill parameters. The *R* loess() function was used with presets span = 0.5 and family = 'symmetric' (Fig. 5A, Supplementary Fig. S5B-C, Supplementary Fig. S6A). To account for the logarithmic scale of the $EC_{50}$ and $n$ Hill model parameters, values were regressed after transformation.

We then screened for Hill parameter-tunable PYL1 positions independent of receptor abundance. To this end, LOESS residuals to each Hill parameter were collected and aggregated by PYL1 position. To screen for positions with significantly higher or lower residual enrichment versus all mutations in the receptor, we calculated position-wise Mann-Whitney U tests prior to Benjamini-Hochberg correction for multiple testing (*91*) (Fig. 5B; Supplementary Fig. S6B).

### 2.6 *In silico* protein stability prediction

We predicted Gibbs free energy of receptor folding ($\Delta\Delta G_f$) *in silico*, based on chain A of the Apo-PYL1 crystal structure (3KAY) (*25*) (Fig. 4F). All PYL1 programmed mutants were used as inputs for ThermoMPNN (*68*) with model weights ‘thermoMPNN_default.pt’. FoldX v5.1 (*92*) was employed as follows: we first used the 'RepairPDB' command against the 3KAY crystal structure, followed by a 'PositionScan' of positions 33-209 of the repaired PDB file. DDMut (*93*) $\Delta\Delta G_f$ calculations were done by block submissions of up to 500 PYL1 mutations via the DDMut web server (https://biosig.lab.uq.edu.au/ddmut/). All PYL1 protein stability predictions can be found in Supplementary Table S4.

### 2.7 Protein structure measurements and display

PDB files of Apo-PYL1 (3KAY) (*25*), (+)-ABA bound PYL1 (3JRS) (*24*) and the PYL1-(+)-ABA-ABI1 complex (3KDJ) (*26*) were downloaded from the RCSB. Using the *R* bio3d package v2.4-4 (*94*), we encoded mutant residual-averaged binding scores or Hill parameter residuals to the abundance-regression in the B-factor column.

PYL1 per-residue conservation values were obtained from ConSurf-DB (*95*). Specifically, 3KDJ (*26*) was used as an input crystal structure and a multiple sequence alignment was computed from 300 related protein sequences across the plant kingdom (Fig. 2A).

Using the PyMOL v2.5.3 (*96*) function get_sasa_relative, we calculated the relative solvent-accessible surface area (rSASA) for residues in the heterotrimeric complex (*26*) (Fig. 2A, Fig. 4C). PYL1 sites with rSASA values of less than or equal to 25% were classified as core residues (Fig. 4D).

Annotated PYL1-(+)-ABA and PYL1-ABI1 protein-protein interface contacts were adopted as by the original analyses of Miyazono *et al.* (*24*) (Fig. 2A), and PYL1-PYL1 interface contacts were annotated by use of GetContacts (https://getcontacts.github.io/; get_static_contacts.py) with the homodimer input structure (*25*) (Fig. 4C). Distances between both PYL1 and hormone, as well as PYL1 and ABI1 heavy atoms were calculated by use of the *R* bio3d package v2.4-4 (*94*) (Fig. 3D, Fig. 5B; Supplementary Table S5).

All visual displays of protein 3D structures were performed using ChimeraX v1.9 (*97*).

### 2.8 Exponential distance decay of PYL1 mutant $EC_{50}$ changes

Mutant absolute $EC_{50}$ fold changes over euclidean distance away from the (+)-ABA or ABI1 interfaces (Fig. 3D) were modelled by an exponential decay process with $= 0$ . Allosteric decay rates (-b) and X-intercept (a) parameterisations were obtained by a nonlinear least squares regression on the log-transformed $EC_{50}$ fold changes:

$$|log_{10}(EC_{50}\, fold\ change)| \approx a \cdot e^{-b \cdot distance}$$

To this end, we used the *R* nls() function with shared initialisations on a = 4 and b = 0.5. 95% confidence intervals were calculated by bootstrapping the model parameter estimates with 1,000 replicates.

### 3. Supplementary Figures

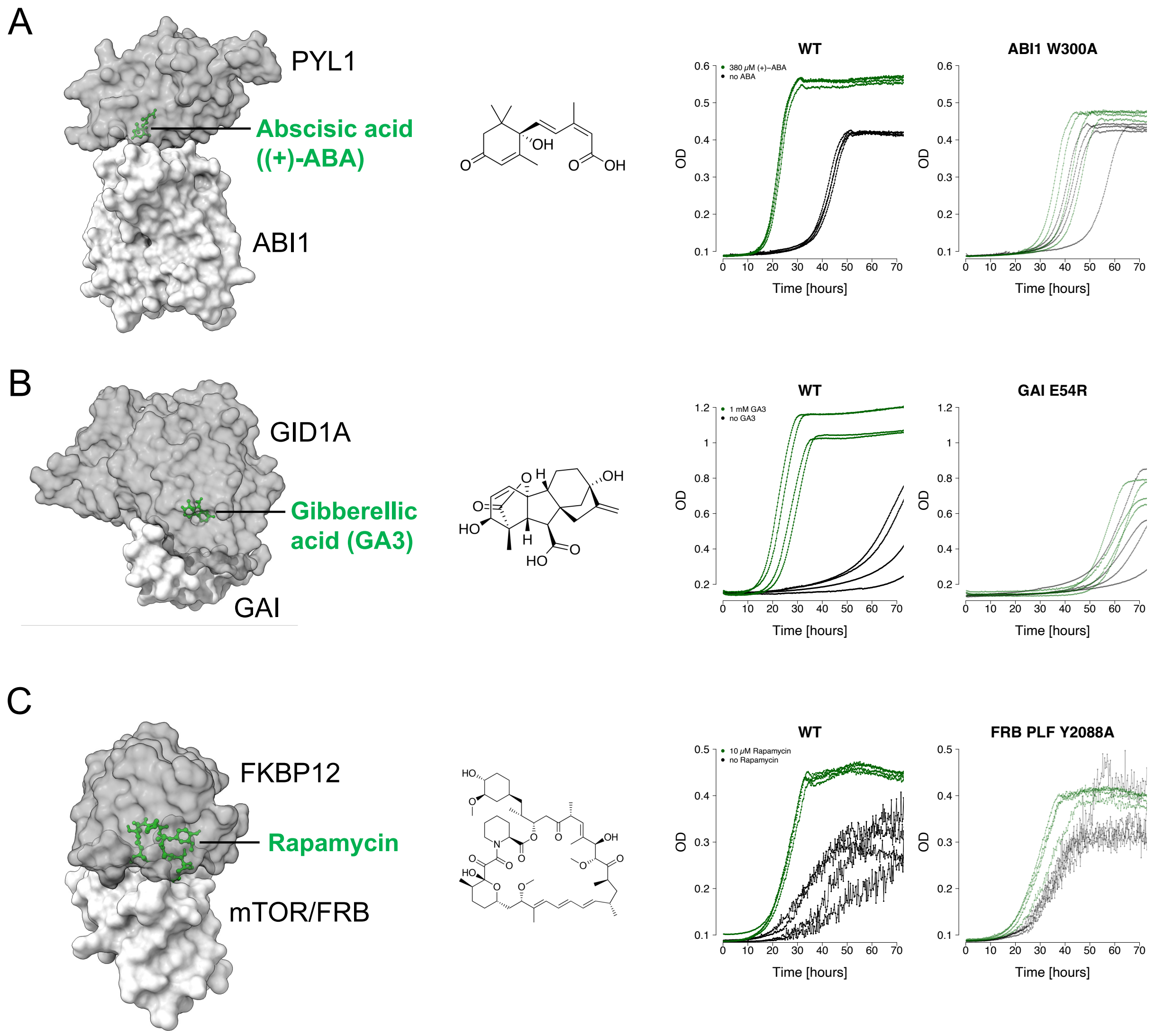

**Supplementary Figure S1: GluePCA scales to diverse chemically-induced dimerisation systems**.
**A:** PYL1-ABI1 binding, mediated by abscisic acid ((+)-ABA). Left: complex structure (PDB: 3KDJ), middle: (+)-ABA molecular structure ($C_{15}H_{20}O_4$; 264.3 g·mol$^{-1}$), right: 70h microtiter plate-based BY4741 yeast colony liquid growth after transformation of the corresponding vector pMS-GluePCA1 (see Supplementary Table S1). Either the wildtype PYL1 and ABI1 proteins (left) or the wildtype PYL1 and mutant ABI1 W300A proteins (right; Miyazono *et al.*, Nature 2009) were inoculated separately. Cells were grown in the presence of 200 µg/mL MTX, enriched with 380 µM (green lines) or 0 µM (black lines) (+)-ABA. Four replicates were tested per condition.
**B:** GID1A-GAI binding, mediated by gibberellic acid (GA3). Left: complex structure (PDB: 2ZSH), middle: GA3 molecular structure ($C_{19}H_{22}O_6$; 346.4 g·mol$^{-1}$), right: 70h microtiter plate-based BY4741 yeast colony liquid growth after transformation of the corresponding vector pMS-GluePCA2 (see Supplementary Table S1). Either the wildtype GID1A and GAI proteins (left) or the wildtype GID1A and mutant GAI E54R proteins (right; Murase *et al.*, Nature 2008) were inoculated separately. Cells were grown in the presence of 200

µg/mL MTX, enriched with 1 mM (green lines) or 0 µM (black lines) GA3. Four replicates were tested per condition.

**C:** FKBP12-mTOR/FRB domain binding, mediated by rapamycin. Left: complex structure (PDB: 1FAP), middle: rapamycin molecular structure ($C_{51}H_{79}NO_{13}$; 914.2 g·mol$^{-1}$), right: 70h microtiter plate-based *MTOR1-1* ∆*fpr1* yeast colony liquid growth after transformation of the corresponding vector pMS-GluePCA3 (see Supplementary Table S1). Either the wildtype FKBP12 and mTOR/FRB proteins (left) or the wildtype FKBP12 and mutant mTOR/FRB PLF Y2088A proteins (right; Stankunas *et al.*, ChemBioChem 2007) were inoculated separately. Cells were grown in the presence of 200 µg/mL MTX, enriched with 10 µM (green lines) or 0 µM (black lines) rapamycin. Four replicates were tested per condition. We note that this rapamycin-resistant yeast strain displays a flocculation phenotype at lower fitness, in line with the observed condition-specific increase in measurement uncertainty.

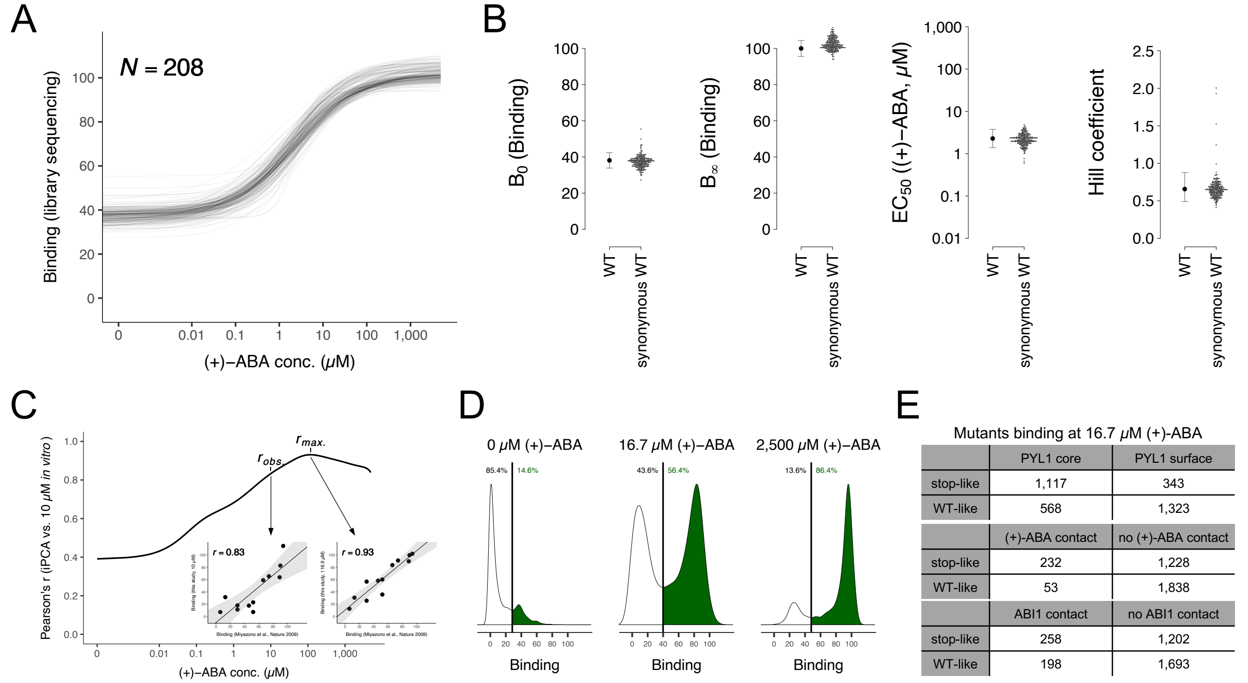

**Supplementary Figure S2: Benchmarking of GluePCA-based deep mutational scanning of PYL1**.

**A:** GluePCA dose-response curves of all 208 synonymous wildtype PYL1 variants.

**B:** Distributions of the four Hill parameters $B_0$, $B_\infty$, $EC_{50}$ and $n$, respectively between the chosen wildtype (left) and the synonymous wildtype variants (right). Error bars indicate 95% confidence intervals.

**C:** Pearson correlation of GluePCA data to PYL1-ABI1 pull-down measurements of twelve PYL1 alanine mutants at 10 µM (+)-ABA (Miyazono *et al.*, Nature 2009). The correlation reaches its maximum at ~117 µM (+)-ABA in GluePCA, suggesting an ~11.7-fold potency shift between yeast GluePCA and *in vitro* studies (see Fig. 1F).

**D:** Kernel density of bulk GluePCA PYL1 mutant library binding fitness at 0, 16.7 and 2,500 µM (+)-ABA. Percentages of wildtype-like binding profiles of PYL1 mutants (green) are assigned based on strict bimodal peak separations (see Fig. 1G).

**E:** Contingency tables of sets of PYL1 mutant binding fitness values at 16.7 µM (+)-ABA (see Fig. 2A).

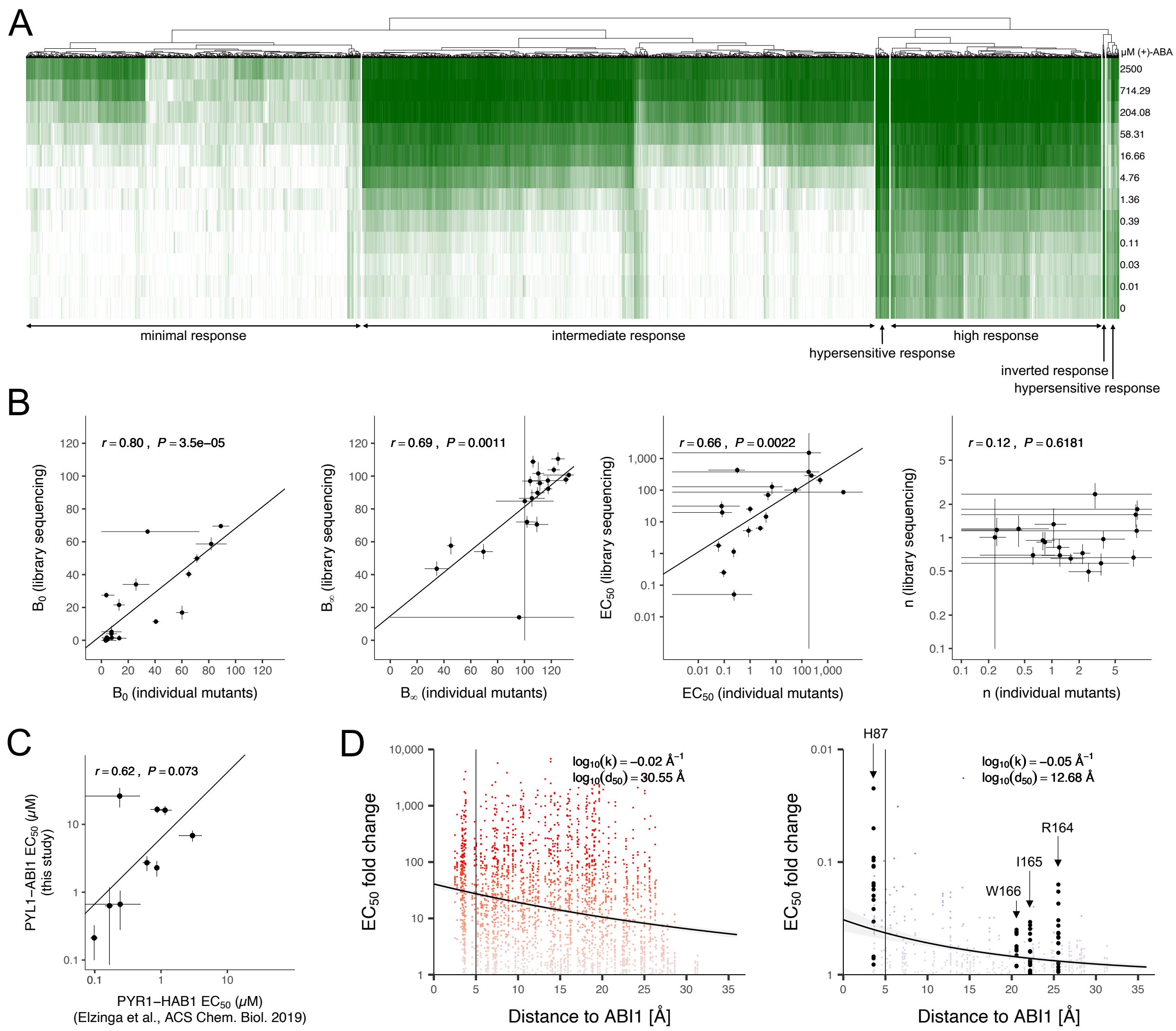

**Supplementary Figure S3: PYL1 mutant categorisation and dose-response parameter validation.**
**A:** Hierarchical clustering of 3,485 PYL1 variants with complete GluePCA measurements across all twelve (+)-ABA concentrations. Binding scores are colour coded in accordance with Fig. 2A.
**B:** Correlation between $B_0$, $B_\infty$, $EC_{50}$ and $n$ Hill parameter estimates of bulk GluePCA PYL1 library mutants and microtiter plate-based dose-response growth measurements of isolated mutants (N = 19; see Materials and Methods). Error bars indicate standard errors of the parameters. Lines represent linear regressions. *r*, Pearson correlation coefficient.
**C:** Correlation between abscisic acid-based $EC_{50}$ estimates of GluePCA PYL1-ABI1 variants and their homologous PYR1-HAB1 variants in a yeast-two-hybrid assay (N = 9; Elzinga *et al.*, ACS ChemBio 2019). Error bars indicate standard errors of the parameters. Line represents linear regression. *r*, Pearson correlation coefficient.
**D:** Euclidean distance decay scatterplot of $EC_{50}$ fold changes within the PYL1 structure. X-axes indicate measured distances between receptor positions side-chain heavy atoms and the ABI1 side-chain heavy atoms. Y-axes indicate absolute $EC_{50}$ fold changes, in order to directly compare the scales of increases (left panel with red-scaled data points) and decreases (right panel with blue-scaled data points). Bold black data points indicate particular sites of interest. Black curves depict regressions of inferred exponential decay, grey shadings indicate the 95% confidence intervals. Vertical black lines at X = 5 Å depict approximate contacts to ABI1. Key rate parameter estimates are shown for both regressions.

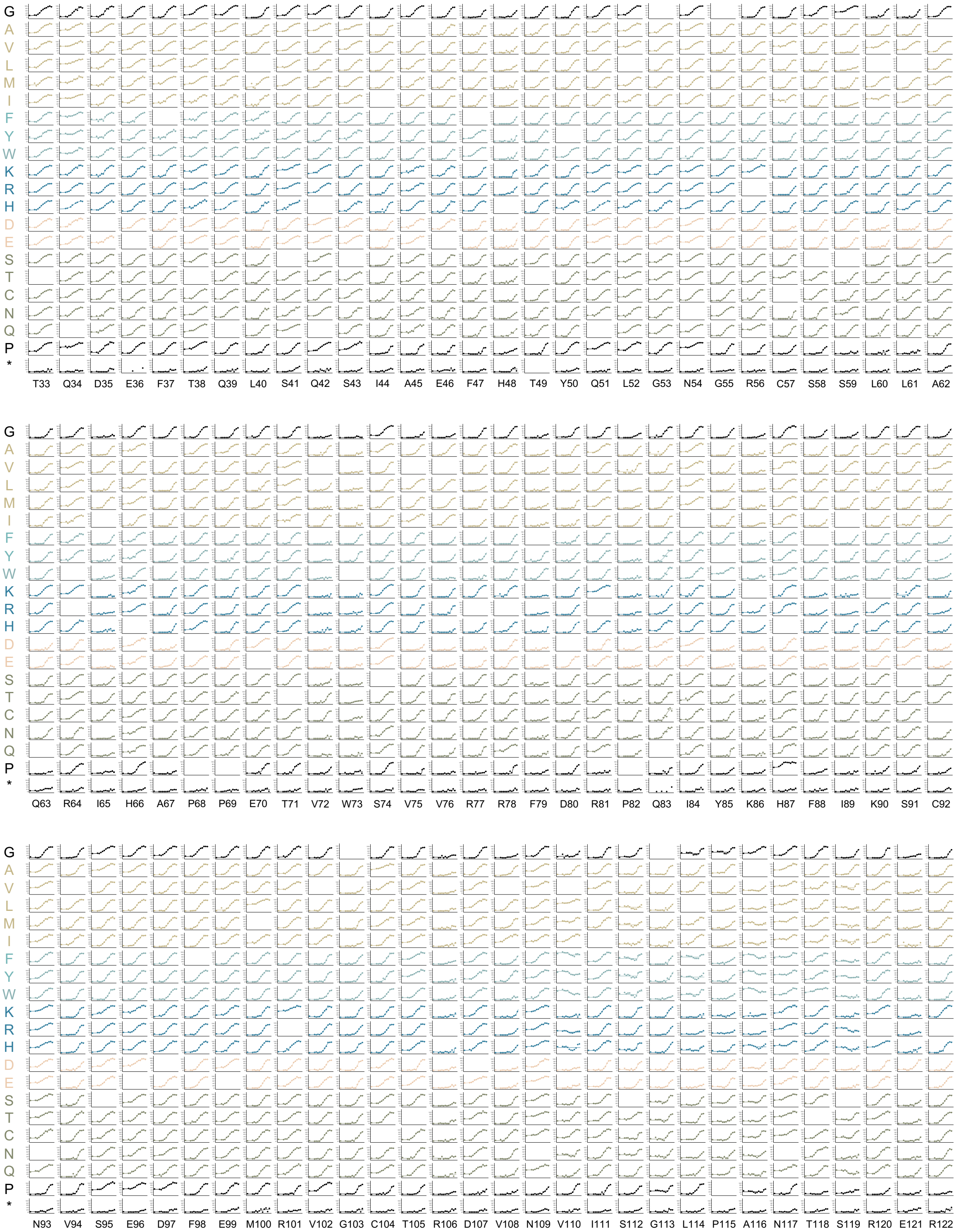
G A V L M I F Y W K R H D E S T C N Q P *
T33 Q34 D35 E36 F37 T38 Q39 L40 S41 Q42 S43 I44 A45 E46 F47 H48 T49 Y50 Q51 L52 G53 N54 G55 R56 C57 S58 S59 L60 L61 A62
G A V L M I F Y W K R H D E S T C N Q P *
Q63 R64 I65 H66 A67 P68 P69 E70 T71 V72 W73 S74 V75 V76 R77 R78 F79 D80 R81 P82 Q83 I84 Y85 K86 H87 F88 I89 K90 S91 C92
G A V L M I F Y W K R H D E S T C N Q P *
N93 V94 S95 E96 D97 F98 E99 M100 R101 V102 G103 C104 T105 R106 D107 V108 N109 V110 I111 S112 G113 L114 P115 A116 N117 T118 S119 R120 E121 R122

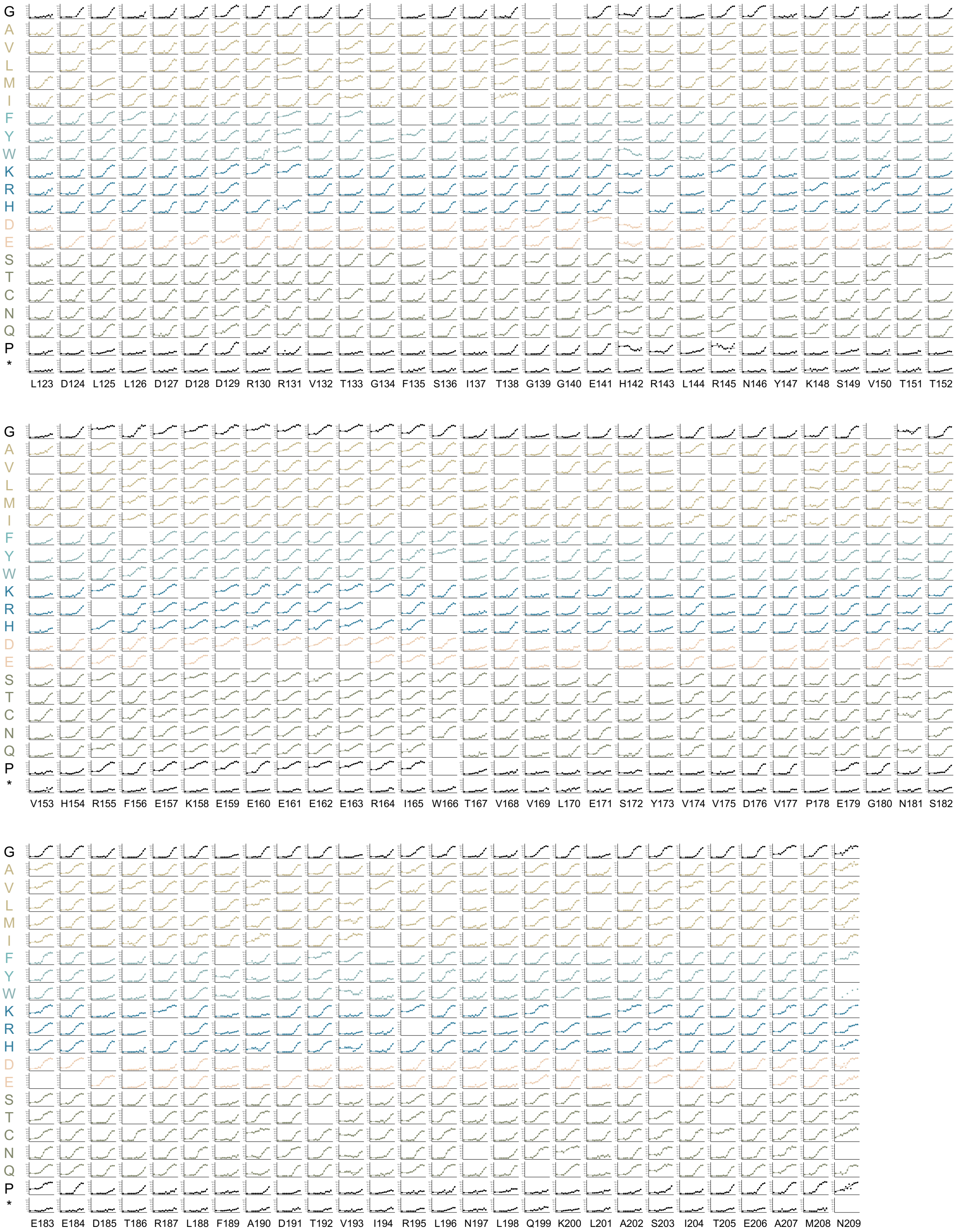

**Supplementary Figure S4: Complete set of 3,528 PYL1 mutant (+)-ABA dose-response curves.**

ABI1-binding dose-response heatmaps of PYL1 mutants exposed to twelve (+)-ABA concentrations. Heatmaps feature the respective PYL1 receptor amino acid sequence along the X-axis, with columns showing the PYL1-ABI1 binding dose-response curves and raw binding scores of all 20 mutant genotypes per residue. Mutant colouring is in accordance with amino acid chemotypes (see Fig. 3F-G). Wildtype variant positions are left blank for improved interpretability.

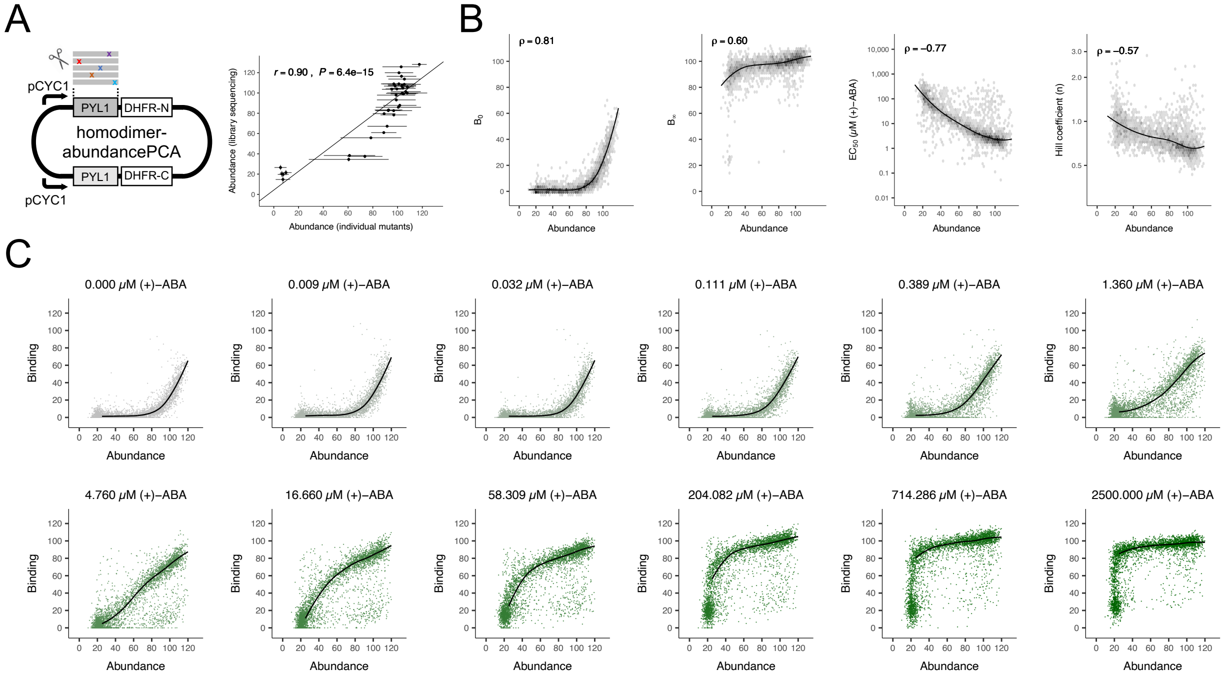

**Supplementary Figure S5: PYL1 mutant homodimer abundance and regressions to Hill parameters.**
**A:** Left: PYL1-PYL1 homodimer PCA setup for deep mutational scanning measurements of PYL1 abundance. On the same plasmid, two CYC promoters express PYL1 mutants as N-terminal fusions to DHFR-N and a sequence-recoded wildtype PYL1 as an N-terminal fusion to DHFR-C. Upon variant library transformation into *S.cerevisiae*, changes in relative cell fractions are obtained through variant competition experiments in which vital DHFR reconstitution is mediated through PYL1-PYL1 association. Right: Correlation between homodimer abundancePCA scores of bulk GluePCA PYL1 library mutants and microtiter plate-based growth measurements of isolated mutants (N = 39; see Materials and Methods). Vertical error bars indicate standard deviations of library-level abundance scores, horizontal error bars depict standard deviations from eight independent yeast colonies per variant in the microtiter plate-based growth assay. Line represents linear regression. *r*, Pearson correlation coefficient.
**B:** Scatterplots of mutant PYL1 homodimer abundance (X-axes) versus the inferred Hill parameters of PYL1-ABI1 binding (Y-axes). Black lines indicate LOESS regressions, grey shading levels scaled by mutant density in each segment. *ρ*, Spearman's rank coefficient.
**C:** Scatterplots of mutant PYL1 homodimer abundance (X-axes) versus the raw PYL1-ABI1 binding scores (Y-axes) at each of the twelve (+)-ABA concentrations. Black lines indicate LOESS regressions for all data points above 25% abundance, data point shading levels increase from lowest (light grey) to highest (dark green) (+)-ABA concentration.

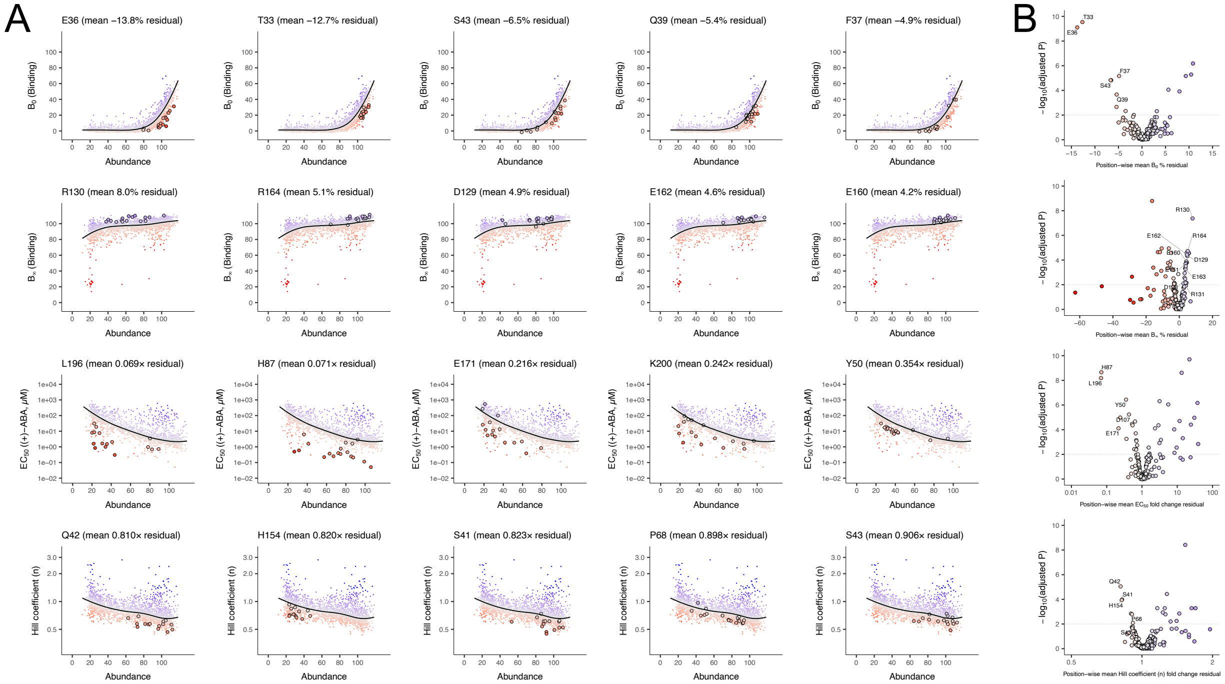

**Supplementary Figure S6: PYL1 positions with significant receptor tunability after abundance normalisation.**

**A:** Scatterplots of mutant PYL1 abundance versus the four different Hill parameters of PYL1-ABI1 binding, with subpanels ordered from top to bottom: $B_0$, $B_\infty$, $EC_{50}$, $n$. Five residue examples of significant tunability were chosen in each case (see Fig. 5). Black lines in each subpanel indicate LOESS regressions, while data points represent individual mutants. Data points are colour scaled by their residual distances to the regression, from low (red) to high (blue). Encircled data points highlight mutations in the chosen example residues of PYL1 which are amenable to Hill parameter tuning independent of protein abundance.

**B:** Volcano plots, displaying inferred position-averaged residual distances to the four Hill parameters' LOESS regressions on the X-axes against mutants' inverse logarithmic *P* value (Mann-Whitney U test) on the Y-axes, adjusted for multiple testing by false-discovery rate (FDR) control. Colour scaling from red to blue is linear along the X-axes. Dashed horizontal lines depict FDR < 0.01. Variants in (A) are highlighted.
